# Absence of Mother’s Curse for performance traits among divergent mtDNAs in heterozygous nuclear backgrounds in Drosophila

**DOI:** 10.64898/2025.12.17.694680

**Authors:** David M. Rand, Faye A. Lemieux, Kenneth M. Bradley, Lindsay Marmor, Leah J. Darwin, Yevgeniy Raynes

## Abstract

Maternal inheritance allows selection to act on mtDNA-encoded effects in females but prevents direct selection on mtDNA in males. Mutations that are deleterious in males but neutral or beneficial in females can persist in populations. This predicts that mtDNA-based phenotypic variation should be more common among males than among females, a pattern referred to as Mother’s Curse (MC). Most studies of MC place alternative mtDNAs on common homozygous nuclear chromosomal backgrounds, a condition not common in nature. Moreover, it is not known whether MC effects accumulate as mtDNAs acquire nucleotide substitutions between populations or species. We tested the MC hypothesis using mtDNAs from *Drosophila melanogaster* (*OreR*, *Zimbabwe* or *w^1118^*), *D. simulans* (*siI* and *siII*) and *D. yakuba* each placed on several *D. melanogaster* nuclear backgrounds heterozygous for different chromosomal deficiencies paired with a common *w^1118^* chromosome set. Females and males were tested for starvation resistance, climbing speed, and flight performance. In the majority of chromosomal backgrounds the variance among mtDNA genotypes was greater in females than in males, opposite from the central prediction of Mother’s Curse. This suggests that additive and dominance variation across the nuclear genome may provide ‘nuclear blessings’ that can counter the curse of maternally inherited mtDNA.

**Teaser text:** Mother’s Curse (MC) posits that selection on mtDNA should be stronger in females than in males due to maternal inheritance of mtDNA. This predicts that phenotypic variation among mtDNA genotypes should be lower for females and higher for males. There is conflicting experimental evidence for MC. Most studies of MC have used a common, homozygous nuclear background and have not explored the influence of divergent mtDNAs as strong predictors of MC effects. We address both issues by assaying performance traits among mtDNAs of varying levels of divergence on heterozygous backgrounds. The data fail to support the MC hypothesis and even reveal the opposite effect that females have greater phenotypic variation across mtDNAs. MC may operate in some contexts, but it is not a consistent force in evolutionary genetics.

## Introduction

Maternal inheritance allows selection to act on mtDNA-encoded effects in females but prevents direct selection on mtDNA in males. Mutations that are deleterious in males but neutral or beneficial in females can persist in populations. This predicts that mtDNA-based disease or phenotypic variation should be more common in males than in females, as haploid selection in females should purge female-specific mtDNA-based variation. This example of genetic conflict was originally introduced by (Cosmides and Tooby 1981) and later gained credibility from differences in the incidence or severity of various disease conditions between males and females (Frank and Hurst 1996). These concepts were reframed as the ‘Mother’s Curse’ by (Gemmell et al. 2004) with relevance to conservation biology concerns about risks of population extinction.

Cytoplasmic male sterility (CMS) in plants is a well-known example of the processes associated with the Mother’s Curse (hereafter, MC) concept. Mutations in mtDNA cause defects in pollen viability with little or no impact on ovules. However, male fertility can be restored by nuclear-encoded restorer loci (e.g., *Rf*, (Melonek et al. 2021) providing clear examples of mitochondrial-nuclear (mitonuclear) interactions. Despite the common occurrence of CMS in plants, examples of CMS or its equivalent in animals are rare (Patel et al. 2016, Dapper et al. 2023). In recent years, a number of studies have reported evidence for the MC pattern of greater phenotypic variation across mtDNA genotypes in males compared to females (Sackton et al. 2003, Innocenti et al. 2011, Camus et al. 2012, Dowling and Adrian 2019). A key assumption in Mother’s Curse is that mtDNA-based phenotypes must be sex-specific with different effects in males and females. The sex-specific differences in phenotypic variation are consistent with a mildly deleterious model of mtDNA mutations. Strong Mother’s Curse scenarios invoke mtDNA mutations that are beneficial in females and deleterious in males, which sweep through populations leading to extinction from male unfitness (Gemmell et al. 2013, Dowling et al. 2015). The sex-specific selective filter that is central to the MC hypothesis has been implicated as a causal factor in faster male aging (Camus et al. 2012), sexual selection (Hill 2018) and models of speciation (Burton and Barreto 2012, Burton 2022). While conceptually appealing (Havird et al. 2019), the generality of these MC effects has been questioned by conflicting data (Mossman et al. 2016, Mossman et al. 2016, Cayuela et al. 2023, Edmands 2024) and evidence from theory (Wade and Brandvain 2009, Wade 2014, Dapper et al. 2023). The MC concept is becoming increasingly common in medical literature as mitochondrial replacement therapy is being practiced in some countries (Reinhardt et al. 2013, Morrow et al. 2015, Eyre-Walker 2017, Hyslop et al. 2025, McFarland et al. 2025). A key question concerning the MC hypothesis is whether mtDNA mutational processes can be significant drivers of phenotypic traits in the face of other sources of phenotypic variation across the genome.

To assess the generality of the MC phenomenon, two technical issues must be addressed that have not been adequately explored in the experimental literature. Most MC analyses use alternative mtDNAs placed on one or more homozygous nuclear chromosomal backgrounds. Since most organisms, including humans, are heterozygous at many loci, we know little about the impact of different mtDNAs on heterozygous, or highly variable nuclear backgrounds.

Second, if the adaptive, sexually antagonistic version of MC is a meaningful evolutionary force, mtDNA data should reveal frequent signatures of positive selection, which is not common (Zwonitzer et al. 2023, Iverson et al. 2025). In mammals and Drosophila, the average dN/dS value for protein-coding genes in mtDNA is lower than that for the nuclear genome, despite much faster divergence for mtDNA at non-coding positions (Saccone et al. 1999, Saccone et al. 2000, Montooth et al. 2009), indicating general patterns of purifying selection. Moreover, we know little about how the depth of mtDNA divergence influences the evidence for or against the patterns predicted by MC.

In the current study we sought to perform experiments that addressed both of these issues. We used mtDNAs that spanned low levels of polymorphism (*Drosophila melanogaster* from different wild-type strains), moderate polymorphism (*D. simulans* strains with *siI* or *siII* mtDNA), to larger degrees of divergence across millions of years between *D. melanogaster, D. simulans,* and *D. yakuba*. These mtDNAs were assayed for starvation resistance, climbing speed, and flight performance on several different heterozygous backgrounds that vary in gene dosage and chromosomal composition. These performance traits were chosen for three reasons: 1) they are important correlates of fitness, 2) they require proper mitochondrial metabolic function, and 3) they are not limited to one sex so direct female-male comparisons can be made. While some experiments produced patterns consistent with the MC hypothesis, across all experiments and nuclear backgrounds the phenotypic variance among mtDNA genotypes was greater in females than in males. This result is the opposite of the Mother’s Curse prediction. Moreover, the impact of the foreign *D. yakuba* mtDNA was similarly neutral or beneficial in both males and females, suggesting long-term stabilizing selection on mtDNA-based performance traits. The mitonuclear epistatic interactions across the different heterozygous backgrounds and different mtDNA haplotypes were also more pronounced in females than males. This suggests that the nuclear variation associated with chromosomally localized gene dosage or dominance effects could overshadow sex-specific selective sieve effects of mtDNAs, or the genomic conflicts, that might generate the patterns of Mother’s Curse.

## Materials and Methods

### Fly strains

Distinct mtDNAs were each placed onto a common *Drosophila melanogaster w^1118^* nuclear genomic background using repeated backcrossing to Bloomington Stock Center stock #6326. Virgin females from each mtDNA line were mated to male *w^1118^*for 10 generations. We used mtDNAs that span low levels of polymorphism in *D. melanogaster* (*w^1118^*, *OregonR* (*Ore*), or *Zimbabwe53* (*Zim53*)), moderate levels of polymorphism in *D. simulans* (*siI* and *siII* (*sm21*)) and deeper divergence within the *D. melanogaster* subgroup (*D. yakuba*). The phylogenetic relationships and numbers of nucleotide and amino acid substitutions among these different mtDNA genomes are shown in Figure 1. Estimates of divergence between strains of *D. melanogaster* are ∼100 to <1000 years, between *D. melanogaster* and *D. simulans* are 1.4-3.4 MYA (Obbard et al. 2012) and between *D. melanogaster* and *D. yakuba* are 8-12 MYA (Tamura et al. 2004, Obbard et al. 2012). The construction of the inter-species mtDNA introgressions has been described elsewhere (Rand et al. 2006, Montooth et al. 2010, Ma and O’Farrell 2015, Spierer et al. 2021). The *D. yakuba* introgression was initially done using microinjection of enriched fraction of *D. yakuba* mitochondria into *D. melanogaster* posterior pole region prior to germ cell cellularization followed by subsequent backcrossing to *D. melanogaster* (Ma et al. 2014, Ma and O’Farrell 2015, Ma and O’Farrell 2016). All mtDNA introgression strains were subsequently backcrossed to *D. melanogaster w^1118^* males as described above, establishing different mtDNAs on a common nuclear genome (Spierer et al. 2021).

**Figure 1.**
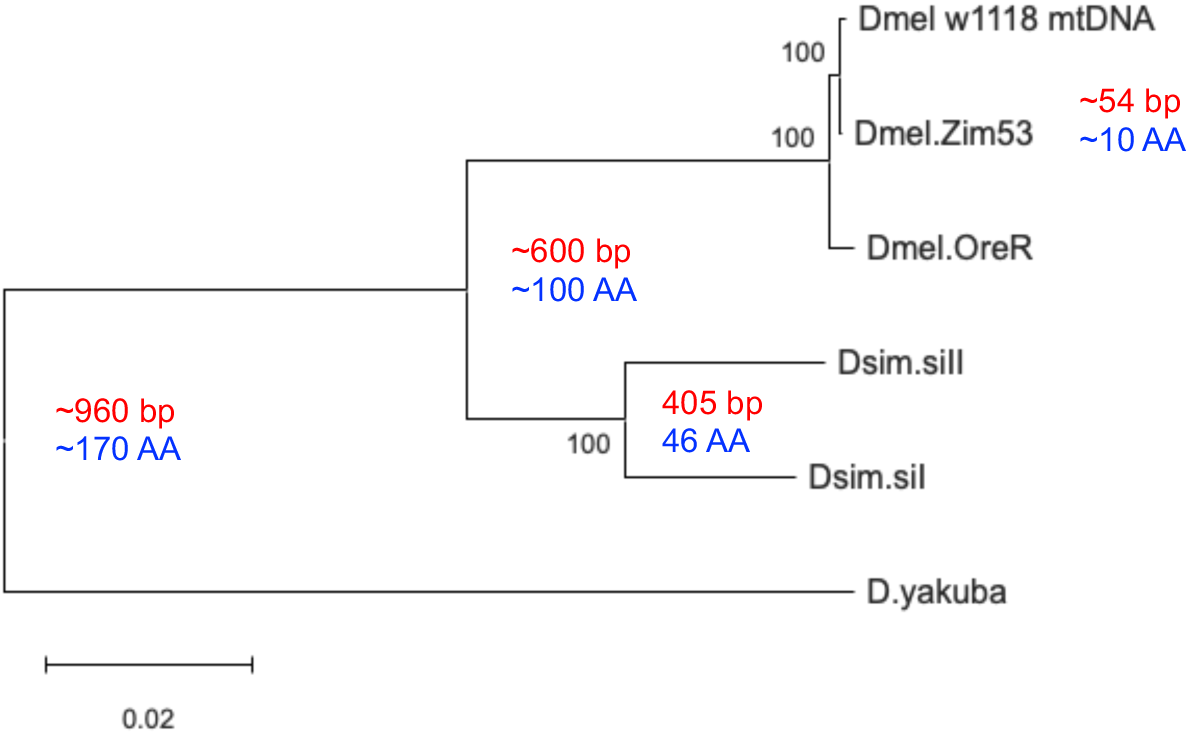
Phylogenetic relationships among the mtDNAs used in these functional assays. A maximum likelihood tree using a Tamura Nei correction model with 500 bootstrap replicates generated in MEGA12. Values inside nodes are the approximate numbers of nucleotide (bp) and amino acid (AA) substitutions between the terminal branches of each clade, based on a parsimony model from (Ballard 2000). Approximate values reflect slight differences in substitutions between different pairs of mtDNAs across clades with more than two mtDNAs. Table 1 shows proportion of differences and Tamura-Nei corrected distances for a multiple alignment of these mtDNAs.

**Table 1.**
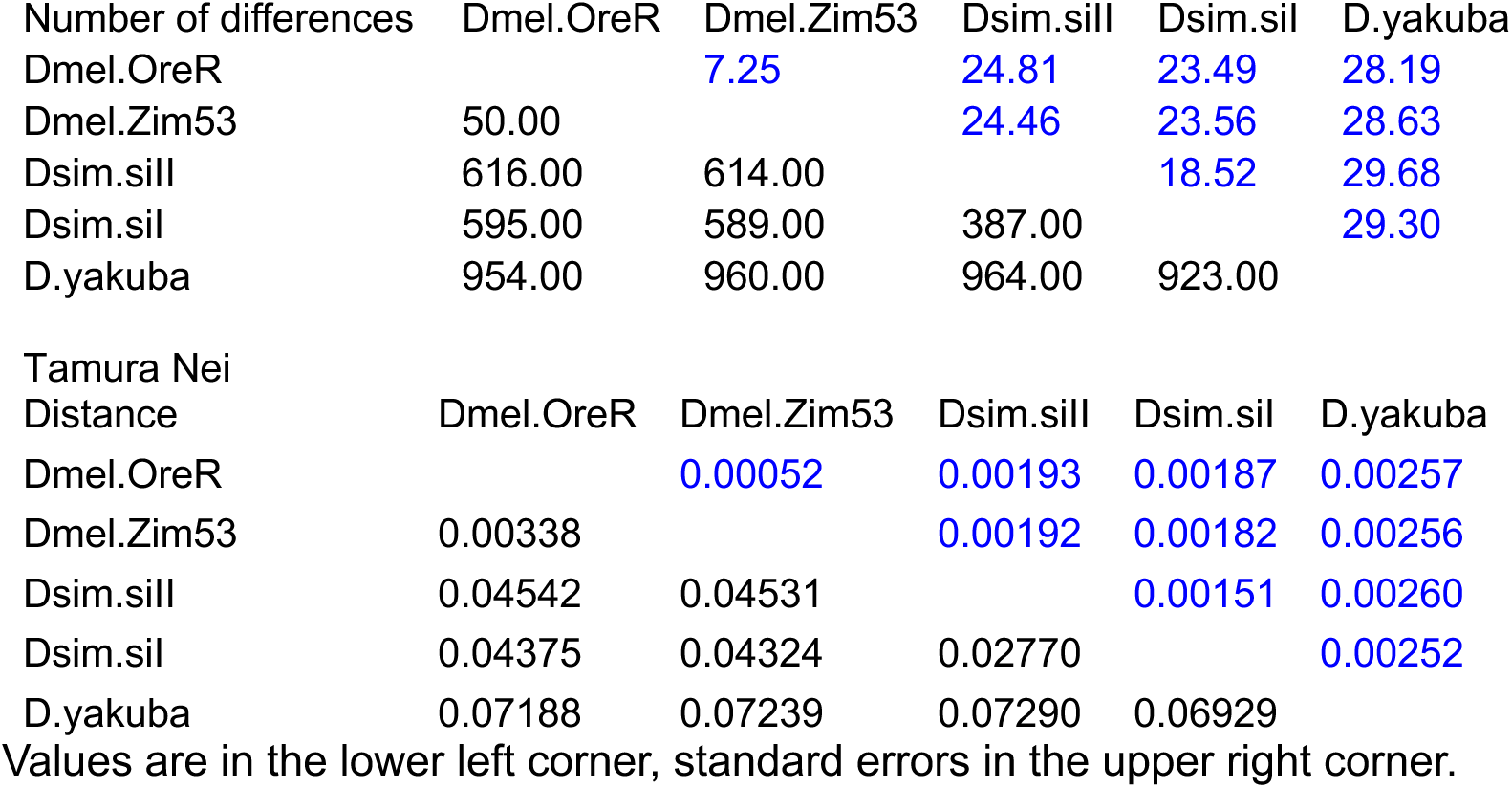
Nucleotide differences and divergence among mtDNAs used in the analyses.

Virgin females of these six mitonuclear lines (notation: mtDNA;nuclearDNA, e.g., *D.sim-siI;w1118* ) were each crossed to males from several different deficiency (Df) stocks that are heterozygous for a 2^nd^ chromosome lacking a single segment of DNA and a second chromosome balancer carrying the curly wing marker CyO (e.g., Df chromosome / SM6 balancer). The experimental flies quantified were the F1 female and male sibling offspring carrying the mtDNA and a *w^1118^* 2^nd^ chromosome from the mother and a deficiency chromosome that lacked the CyO marker from the father. This resulted in each different mtDNA being placed onto a common genome-wide heterozygous background including the shared hemizygous region unique to each Df stock (supplementary Figure S1). Since natural populations have variation in gene copy number and various levels of insertion/deletion polymorphism, this survey of different deficiency stocks provides a model for that kind of nuclear variation, in addition to masking deleterious recessive mutations that might be expressed in a homozygous *w^1118^* background. All Df stocks were obtained from the Bloomington Drosophila Stock Center and are listed in Table S1.

The deficiency regions vary in length and accordingly differ in the number and functions of genes rendered hemizygous in the F1 flies for which phenotypes were quantified (range: 23 – 146 genes; see Table S1). The different Df chromosomes were constructed from *w^1118^* chromosomes but differ slightly in the source of the *w^1118^* stock (Table S1). While the comparisons across mtDNAs within a particular heterozygous Df/*w^1118^*F1 background will have the same chromosome pairs, each specific mtDNA placed onto a different heterozygous Df/*w^1118^*F1 background will differ in the set of hemizygous chromosomal regions and in the origin of the *w^1118^* from which the different Dfs were generated. Due to the number of F1 adults needed to quantify the three performance traits and variation in the number of available parents of each genotype when crosses were conducted, the experiments were done in different blocks. We did not pair all different mtDNAs with all seven Df stocks in each block.

### Phenotypes studied

Three quantitative traits were assayed in sibling females and males across the different blocks of the experiment: climbing speed, flight performance, and starvation resistance. These traits are metabolically demanding of muscle, neuronal and anabolic and catabolic functions that should challenge mitochondrial processes. Moreover, the traits are not sex-limited, i.e., they are shared by both sexes and can be sexually dimorphic, allowing for a fair test of the Mother’s Curse hypothesis (hereafter MCH). For all experiments, fly cultures were density controlled with equal numbers of parents for 2-3 days of egg lay after which adults were allowed to eclose, mature and mate for ∼5 days. Adults were sorted into single sex holding vials on standard Drosophila media.

<u>Climbing speed</u> was quantified using the FreeClimber apparatus and software (Spierer et al. 2021). Climbing videos are recorded for five seconds using a Raspberry Pi camera triggered by a photo-activated switch connected to the climbing rig. The FreeClimber software tracks individual flies and calculates climbing speed as the two-second window of the video during which the value of vertical position vs. time (i.e., slope) is the steepest and most linear. The data are averaged across three successive gentle ‘knockdowns’ of the rack of six replicate vials of 15-20 adults that place the flies at the bottom of each vial and induce climbing or negative geotaxis in the rack of six vials. The slope term for each vial is the mean across the three replicate ‘knockdowns’, and ANOVAs were performed on these means across the six replicate vials in each genotype-sex-age sample based on 6 vials x 15 flies ≥ 900 individuals per sample. The climbing assay was done in three distinct blocks of mitonuclear genotypes each with different nuclear deficiency genotypes. The mtDNAs used in these blocks were shared across the three blocks but some mtDNAs were missing from some blocks. Climbing assays were done at two ages (10 and 22, 10 and 24, and 7 and 24 days for the three different blocks) with each age data set analyzed separately. With three blocks of several mitonuclear genotypes, two sexes and two ages, the data for each block is derived from >10,000 individual adult flies.

<u>Flight performance</u> was measured following the protocol refined by (Babcock and Ganetzky 2014) and reported earlier (Spierer et al. 2021). Each sex and genotype combination consisted of approximately 50-100 flies, divided into groups of 10-20 flies across five glass Drosophila culture vials. After introduction to the flight column, free-falling flies instinctively right themselves, fly laterally some distance before being immobilized at their respective landing height on a sticky mylar sheet lining the inside wall of the column. A digital image of the mylar sheet was recorded on a fixed Raspberry Pi Camera (V2) and the x,y coordinates of all flies were located with the ImageJ/FIJI Find Maxima function. For each sex-genotype combination, the mean landing height was calculated for only the flies that landed on the acrylic sheet. As a sacrificial assay, flight was quantified at one age.

<u>Starvation resistance</u> was measured in single-sex assays by placing approximately 100 flies per genotype in each of three replicate demography cages. Flies were collected and sorted over CO_2_ and allowed to recover in quart-sized plastic demography cages affixed with a 25×95 mm vial containing standard Drosophila food media. The three replicate cages for each genotype were allowed to recover for one day, any dead flies were removed, and the vial for each demography cage was replaced with 2% Difco Bacto-agar prepared with deionized H_2_0 to initiation starvation. The number of dead flies was recorded every 12 hours and removed from each demography cage until all flies were dead. The survival data from each cage were generally consistent so counts from each replicate cage were pooled and the combined data modeled using a Kaplan-Meier survival model implemented in R (Therneau and Grambsch 2000) and confirmed using JMP Pro® version 17. Significant differences in survivals were determined using a log rank test.

<u>Tests of the Mother’s Curse Hypothesis (MCH).</u> All data sets were quantified as factorial ANOVAs with mtDNA genotype and nuclear genotype as main effects plus the mtDNA x nuclear interaction term, using the R functions aov or lm (R_Core_Team 2021). The climbing assay was done at two ages, so age was a third main effect in those models. To test the MCH, we calculated the mtDNA-based genetic coefficient of variation (CVmt) for males and females separately. For each mtDNA in each nuclear background a mean value of climbing, flight or starvation was determined for each sex and age. The standard deviation of these mean values across mtDNAs was divided by the mean value across these mtDNAs to provide the CVmt values. The MCH is supported if CVmt-males > CVmt-females; the MC hypothesis is not supported if CVmt-males = CVmt-females, and the MC hypothesis is falsified if CVmt-males < CVmt-females. The number of CVmt values within each block is rather small (3-8 values per sex), so tests within each block are underpowered, however the collective outcomes of all assays can be tested with a one-way ANOVA or a Wilcoxon signed rank test. To visualize these tests, biplots were generated with the CVmt values in each nuclear background plotted on the X- and Y values for females and males, respectively.

## Results

### mtDNA divergence

The complete sequences of the five different mtDNAs have been published and analyses of nucleotide, amino acid, and RNA divergence indicate the extent of functional and, presumably, neutral divergence that has accumulated (Ballard 2000, Ballard 2000, Montooth et al. 2009). Figure 1 and Table 1 show considerable nucleotide divergence between strains and species as is well known for mtDNA. A notable feature of amino acid differences is a strong pattern of purifying selection on most mtDNA-encoded proteins. Average amino acid divergence among mtDNA protein-coding genes is lower than the average divergence among nuclear-encoded proteins. This indicates that the strong mutation pressure, implied by high synonymous divergence, results in considerable purging of deleterious amino acid mutations that slows overall divergence (Montooth et al. 2009). The molecular divergence suggests that mtDNAs from other species likely carry amino acid and RNA mutations that are less well adapted to the nuclear-encoded proteins with which they interact. These mutations are candidates for the phenotypic effects predicted by the MC hypothesis.

### Climbing speed, block 1

Figure 2 shows the results for climbing speed assay in females and males where four different mtDNAs were placed on four different nuclear backgrounds at two ages (10 and 22 days). Three backgrounds are heterozygous for a deficiency chromosome (1491, 4959, 7837) and the *w^1118^* chromosome, and the fourth background is the homozygous *w^1118^* chromosome set used as a reference genotype for each experiment. The *w^1118^* mtDNA paired with the Df_7837 and the *w^1118^* nuclear backgrounds were not quantified. Data are based on six replicate vials of 15-20 flies per vial for each mitonuclear genotype, each assayed in three replicate drop-induced climbing videos, totaling >13,000 flies. The ANOVAs show significant variation among mtDNAs, nuclear backgrounds and their interaction (Table 2). For females, Df_1491 reduces climbing speed in all mtDNAs, while this background results in average performance in males. The different nuclear backgrounds have similar effects on climbing speed for flies of different ages, but the patterns across females (Figure 2A,B) and males (Figures 2C,D) are distinct. Tables 2A,B show that for females, the nuclear F-value is less than the mtDNA F-value, while the opposite is true for males. The effect of age (‘day’ in the tables) is not prominent in either females or males. Notably the variation among mtDNAs is higher in females than males (Table 2, F = 29.38 for females, F = 3.88 for males).

**Figure 2.**
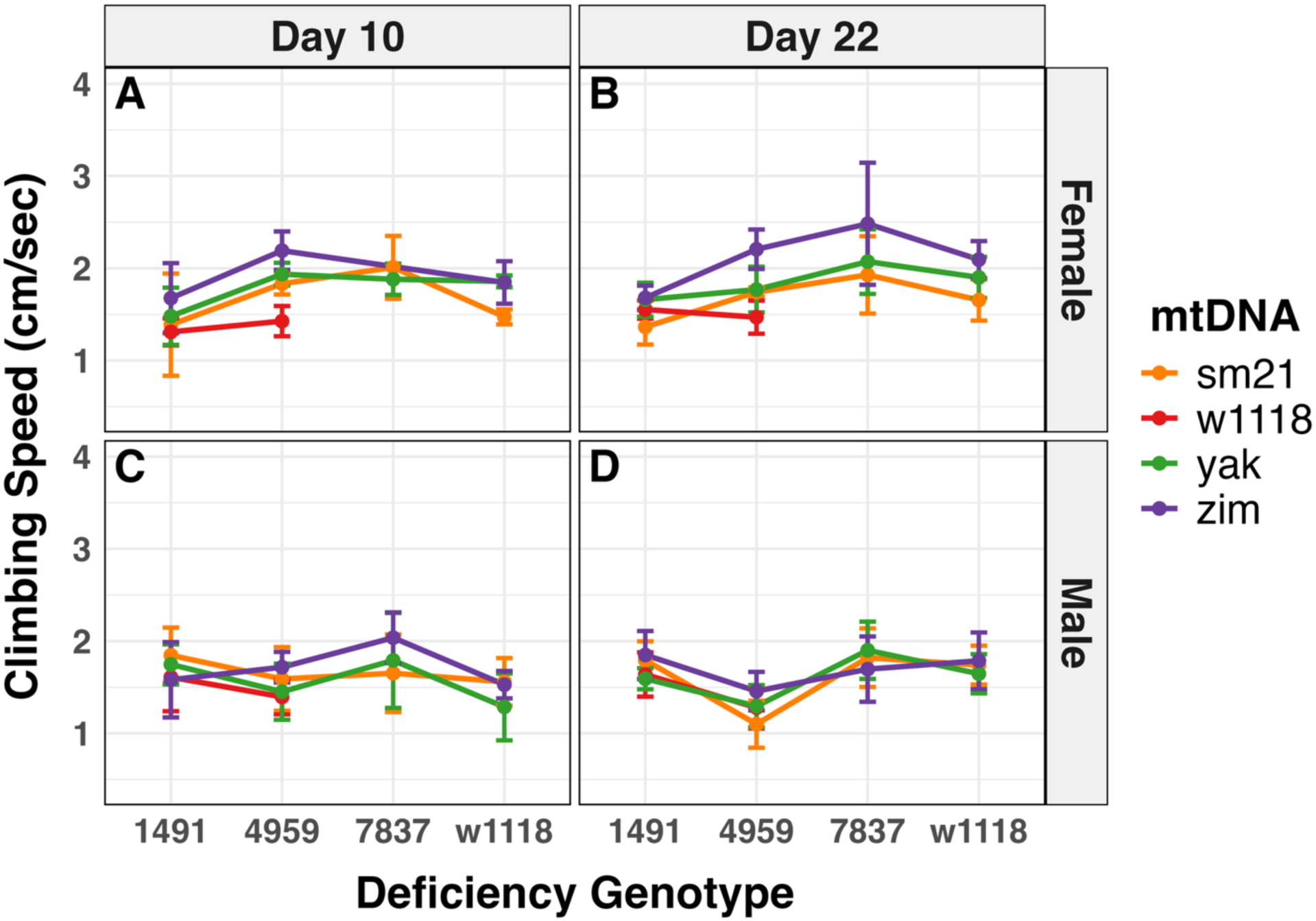
Climbing Speed, block 1. Climbing speed was quantified for four mtDNAs paired with 4 nuclear backgrounds at two ages (10 and 22 days) in females and males (Figure 2A, B, C, D, respectively). There are significant mtDNA, nuclear and mitonuclear interaction effects in males and females. The variation among mtDNAs (F-value) is greater in females than in males. The w1118 mtDNA was not quantified in the 7837 and *w^1118^* nuclear backgrounds, but ANOVA results are qualitatively the same (See Table 2 and Table S1). Sample sizes exceed 10,000 adult flies; see Methods.

**Table 2A,B.** Analysis of variation for climbing speed in block 1 among 16 mitonuclear genotypes of females and males.

| FEMALES | Df | Sum Sq | Mean Sq | F value | Pr(>F) |  |
| --- | --- | --- | --- | --- | --- | --- |
| mito | 3 | 5.901 | 1.9670 | 29.380 | 1.10e-14 | *** |
| geno | 3 | 5.239 | 1.7464 | 26.084 | 2.26e-13 | *** |
| day | 1 | 0.198 | 0.1979 | 2.956 | 0.08784 | . |
| mito:geno | 7 | 1.305 | 0.1865 | 2.786 | 0.00974 | ** |
| mito:day | 3 | 0.097 | 0.0324 | 0.483 | 0.69449 |  |
| geno:day | 3 | 0.318 | 0.1059 | 1.582 | 0.19667 |  |
| mito:geno:day | 6 | 0.252 | 0.0420 | 0.628 | 0.70761 |  |
| Residuals | 135 | 9.038 | 0.0670 |  |  |  |

| MALES | Df | Sum Sq | Mean Sq | F value | Pr(>F) |  |
| --- | --- | --- | --- | --- | --- | --- |
| mito | 3 | 0.888 | 0.2962 | 3.883 | 0.010547 | * |
| geno | 3 | 3.530 | 1.1766 | 15.426 | 1e-08 | *** |
| day | 1 | 0.008 | 0.0083 | 0.109 | 0.741371 |  |
| mito:geno | 7 | 0.625 | 0.0893 | 1.171 | 0.323239 |  |
| mito:day | 3 | 0.054 | 0.0181 | 0.237 | 0.870647 |  |
| geno:day | 3 | 1.466 | 0.4887 | 6.407 | 0.000424 | *** |
| mito:geno:day | 7 | 0.970 | 0.1386 | 1.818 | 0.088348 | . |
| Residuals | 140 | 10.678 | 0.0763 |  |  |  |

Signif. codes: 0 '\*\*\*' 0.001 '\*\*' 0.01 '\*' 0.05 '.' 0.1 ' ' 1 'mito' = mtDNA genotype, 'geno' = nuclear deficiency genotype, 'day' = age

We performed the analyses on a reduced data set of 12 mitonuclear genotypes in which the *w^1118^* mtDNA was removed from all nuclear backgrounds providing a full factorial comparison (3 mtDNAs x 4 nuclear backgrounds). This smaller data set reduces the mtDNA F-value for females compared to the results for 16 mitonuclear genotypes, but this value remains substantially larger than the mtDNA F-value for males. In general, the ANOVAs are qualitatively similar with small differences in the interaction plots and ANOVA table values (compare Figures 2A-D with supplementary Figure S3A-D and Tables 2A,B with Tables 2C,D).

**Table 2C,D.** Analysis of variation for climbing speed in block 1 among 12 mitonuclear genotypes of females and males.

| FEMALES | Df | Sum Sq | Mean Sq | F value | Pr(>F) |  |
| --- | --- | --- | --- | --- | --- | --- |
| mito | 2 | 2.767 | 1.3833 | 18.439 | 1.13e-07 | *** |
| geno | 3 | 5.913 | 1.9711 | 26.274 | 5.12e-13 | *** |
| day | 1 | 0.113 | 0.1127 | 1.503 | 0.223 |  |
| mito:geno | 6 | 0.641 | 0.1068 | 1.423 | 0.212 |  |
| mito:day | 2 | 0.049 | 0.0245 | 0.326 | 0.723 |  |
| geno:day | 3 | 0.263 | 0.0876 | 1.168 | 0.325 |  |
| mito:geno:day | 5 | 0.247 | 0.0495 | 0.660 | 0.655 |  |
| Residuals | 115 | 8.628 | 0.0750 |  |  |  |

| MALES | Df | Sum Sq | Mean Sq | F value | Pr(>F) |  |
| --- | --- | --- | --- | --- | --- | --- |
| mito | 2 | 0.337 | 0.1685 | 2.150 | 0.120895 |  |
| geno | 3 | 3.038 | 1.0127 | 12.927 | 2.25e-07 | *** |
| day | 1 | 0.003 | 0.0030 | 0.038 | 0.845028 |  |
| mito:geno | 6 | 0.624 | 0.1041 | 1.328 | 0.249750 |  |
| mito:day | 2 | 0.048 | 0.0240 | 0.306 | 0.736806 |  |
| geno:day | 3 | 1.469 | 0.4897 | 6.251 | 0.000559 | *** |
| mito:geno:day | 6 | 0.935 | 0.1559 | 1.990 | 0.072253 | . |
| Residuals | 120 | 9.401 | 0.0783 |  |  |  |

Signif. codes: 0 '\*\*\*' 0.001 '\*\*' 0.01 '\*' 0.05 '.' 0.1 ' ' 1 'mito' = mtDNA genotype, 'geno' = nuclear deficiency genotype, 'day' = age

The Mother’s Curse (MC) hypothesis predicts that the phenotypic effects of mtDNA in males will be greater than that in females. To test this, the coefficients of variation for climbing speed across mtDNA genotypes (CVmt) in males and females are plotted against one another (Figure 3A). A CVmt value is calculated within each nuclear genotype x age category separately for females (x-axis, CV_female_) and males (y-axis, CV_male_). The majority of values fall below the 1:1 line indicating the mtDNA genotype effects are greater in females than males, opposite to the prediction of the MC hypothesis. Figure 3B shows the same plot using the reduced data set (shown in Figure S3A-D), with a similar result. To examine the effect of variation in the mean climbing speed values on the generality of these CVmt biplots, bootstrap analyses were performed on the mean values. The results show that the general patterns are confirmed (supplementary Figure S4A,B).

**Figure 3.**
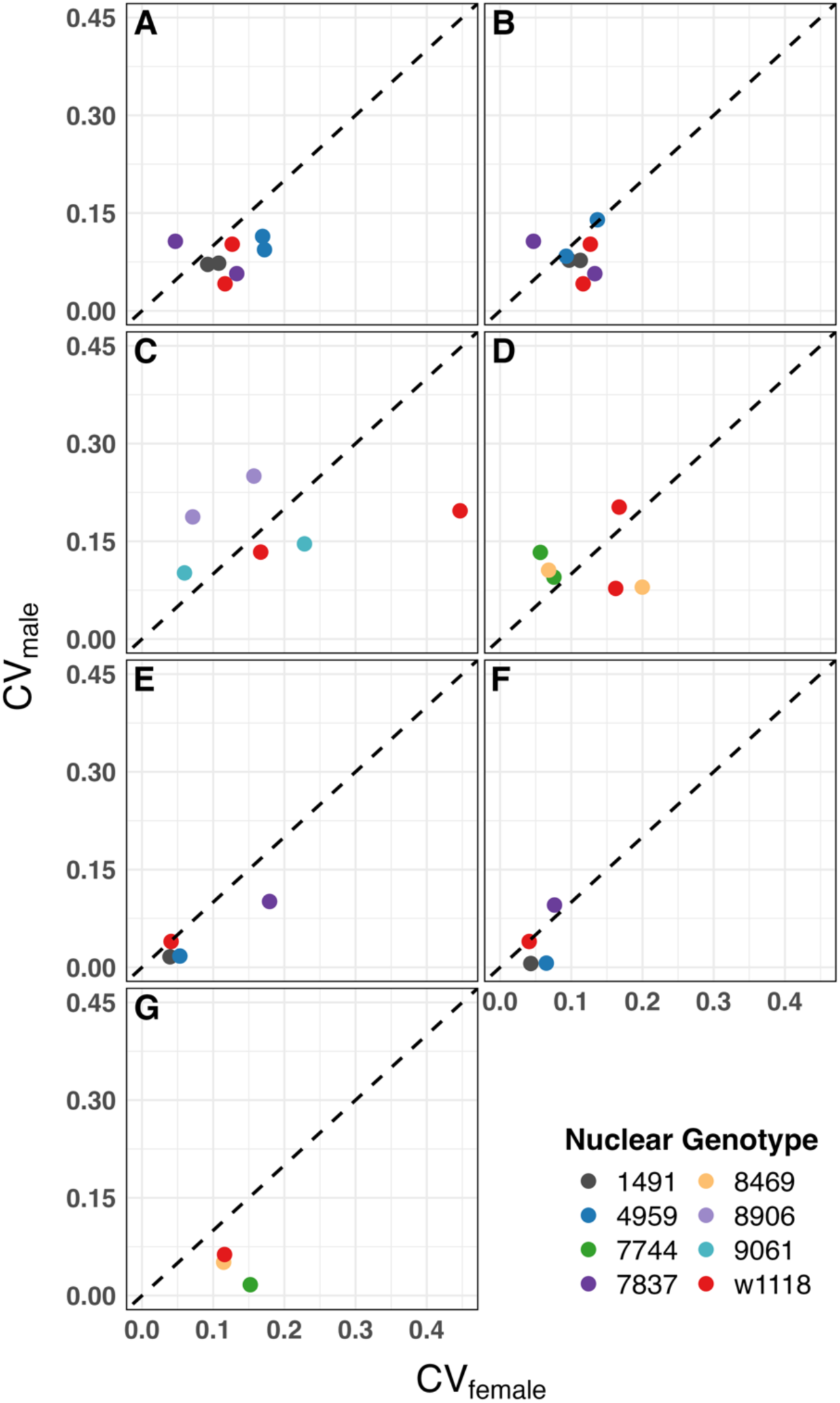
Biplot tests of the Mother’s Curse hypothesis. The coefficients of variation across mtDNA genotypes (CVmt) within each nuclear genotype (or age) category are plotted for females (x-axis, CV_female_) and males (y-axis, CV_male_). A. Climbing speed block 1. B. Climbing speed block1, reduced data set. C. Climbing speed block 2. D. Climbing speed block 3. E. Flight performance, full data set. F. Flight performance using reduced data set with balanced 3×4 design. G. Starvation resistance. Bootstrap analyses of these CVmt values are shown in supplementary figures

We note that all three Df stocks have slightly different chromosomal (genome-wide) backgrounds in addition to the obvious effects of the deficiency (see Table S1). Df_7837 was built from a *w^1118^* stock and is visually distinct in its values from the homozygous *w^1118^* for the various mtDNAs, suggesting a primary role for the set of genes in the deficiency for this effect. This is more apparent at the older age in females (Figure 2B) and the younger age in males (Figure 2C).

### Climbing speed, block 2

Climbing speed was quantified in a second block of 15 mitonuclear genotypes where five mtDNAs (two from *D. melanogaster*, two from *D. simulans* and one from *D. yakuba*) were each placed on two additional deficiencies plus the reference *w^1118^* backgrounds at two ages (7 and 24 days; females: Figure 4A,B; males: Figure 4C,D). The Df_9061 background clearly improved climbing speed for all mtDNAs relative to Df_8906 or the reference *w^1118^*in both females and males (Table 3: nuclear F-value = 32.3 and 65.8, respectively). There was significant variation among mtDNAs across nuclear backgrounds, as well as significant mtDNA x nuclear (mitonuclear) interaction evident by the crossing of lines in these interaction plots. Age had a significant effect in females, but less of an effect in males (Table 3: the F-values for ‘day’ = 93.3 and 5.6 respectively). As in the first block, the effect of mtDNA genotype was greater in females than in males (Table 3: mtDNA F-values = 13.35 and 3.96 for females and males, respectively). The mitonuclear interaction term was significant in both males and females, and the F-value was larger in males. Alternative statistical models have subtle quantitative, and no major qualitative, effects on results (see Table S3).

**Figure 4.**
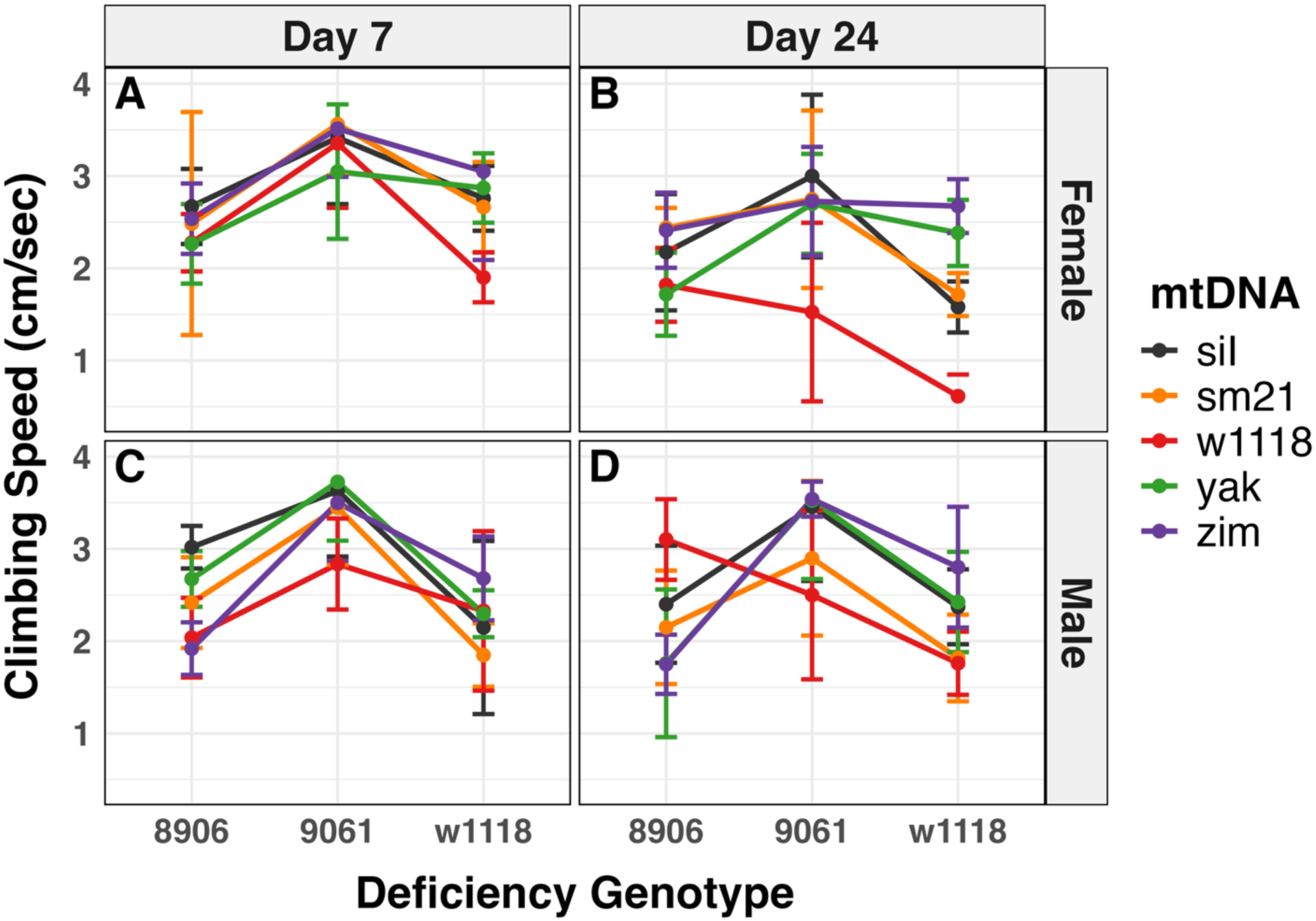
Climbing speed, block 2. Five different mtDNAs paired with two deficiencies and the *w^1118^*background. Significant effects of mtDNA, nuclear genotype and age are revealed in both males and females. The variation across mtDNAs is greater in females than males, in part due to different interactions with age.

**Tables 3A,B.** Analysis of variation for climbing speed in block 2 among 15 mitonuclear genotypes of females and males.

| FEMALES | Df | Sum Sq | Mean Sq | F value | Pr(>F) |  |
| --- | --- | --- | --- | --- | --- | --- |
| mito | 4 | 16.68 | 4.169 | 13.349 | 2.49e-09 | *** |
| geno | 2 | 20.18 | 10.089 | 32.303 | 2.16e-12 | *** |
| day | 1 | 20.57 | 20.573 | 65.869 | 1.61e-13 | *** |
| mito:geno | 8 | 8.99 | 1.123 | 3.597 | 0.000754 | *** |
| mito:day | 4 | 3.41 | 0.852 | 2.729 | 0.031339 | * |
| geno:day | 2 | 2.63 | 1.315 | 4.211 | 0.016618 | * |
| mito:geno:day | 8 | 3.43 | 0.429 | 1.374 | 0.212352 |  |
| Residuals | 150 | 46.85 | 0.312 |  |  |  |

| MALES | Df | Sum Sq | Mean Sq | F value | Pr(>F) |  |
| --- | --- | --- | --- | --- | --- | --- |
| mito | 4 | 5.02 | 1.256 | 3.955 | 0.00442 | ** |
| geno | 2 | 41.79 | 20.896 | 65.807 | < 2e-16 | *** |
| day | 1 | 1.01 | 1.013 | 3.190 | 0.07609 | . |
| mito:geno | 8 | 13.69 | 1.711 | 5.388 | 5.76e-06 | *** |
| mito:day | 4 | 1.03 | 0.257 | 0.808 | 0.52166 |  |
| geno:day | 2 | 0.38 | 0.190 | 0.598 | 0.55137 |  |
| mito:geno:day | 8 | 7.60 | 0.950 | 2.991 | 0.00387 | ** |
| Residuals | 150 | 47.63 | 0.318 |  |  |  |

Signif. codes: 0 '\*\*\*' 0.001 '\*\*' 0.01 '.' 0.1 ' ' 1
'mito' = mtDNA genotype, 'geno' = nuclear deficiency genotype, 'day' = age

The biplot test of the MC hypothesis for climbing speed block 2 is shown in Figure 3C. There are three nuclear backgrounds and two ages resulting in six CVmt pairs of points, roughly equally split on either side of the 1:1 line. This result is not consistent with the MC hypothesis that CVmt-males should be greater than CVmt-females. A bootstrap resampling analysis of the CVmt values is shown in supplementary Figure S5.

The comparison across nuclear backgrounds is more standardized in this block since Df_8906 and Df_9061 were generated from a common *w^1118^* background (Table S1), so the F1 flies share a common chromosomal background and differ only in the loci rendered hemizygous by the deficiencies. The line-crossing in both female and male data for age 24 days (Figure 4B,D) is considerable. It is notable that the *w^1118^* mtDNA is the slowest climber in its own homozygous nuclear background. However, when placed in a genome wide heterozygote for *w^1118^* chromosomes, while also being hemizygous for the genes missing in Df_8906 and Df_9061, that defect in climbing speed appears to be ‘rescued’. These observations indicate that gene dosage, genome wide heterozygosity, and mtDNA variation can all contribute to epistatic interactions influencing climbing.

### Climbing speed, block 3

We quantified climbing speed in a third block of nine mitonuclear genotypes at two ages (10 and 24 days): one mtDNA each from *D. melanogaster*, *D. simulans* and *D. yakuba* was placed on each of two additional deficiencies (Df_7744 and Df_8469) plus the reference *w^1118^* background (females: Figure 5A,B; males: Figure 5C,D). The results show distinct effects in females and males with more pronounced genetic and age effects in males than in females, but more pronounced mitonuclear interaction effects in females (Tables 4A, B). The interaction plots show clear line crossing in females but highly parallel lines in males (Figure 5A-D). The Df_8469 nuclear background increased climbing speed in young and old males (ages 10 and 24 days) and the reference *w^1118^* background showed slower climbing speed, a pattern observed in other experimental blocks for certain mtDNAs paired with the *w^1118^* nuclear background. We note that the reference *w^1118^* background is generally a weak stock and it frequently produced fewer offspring for the experimental crosses.

**Figure 5.**
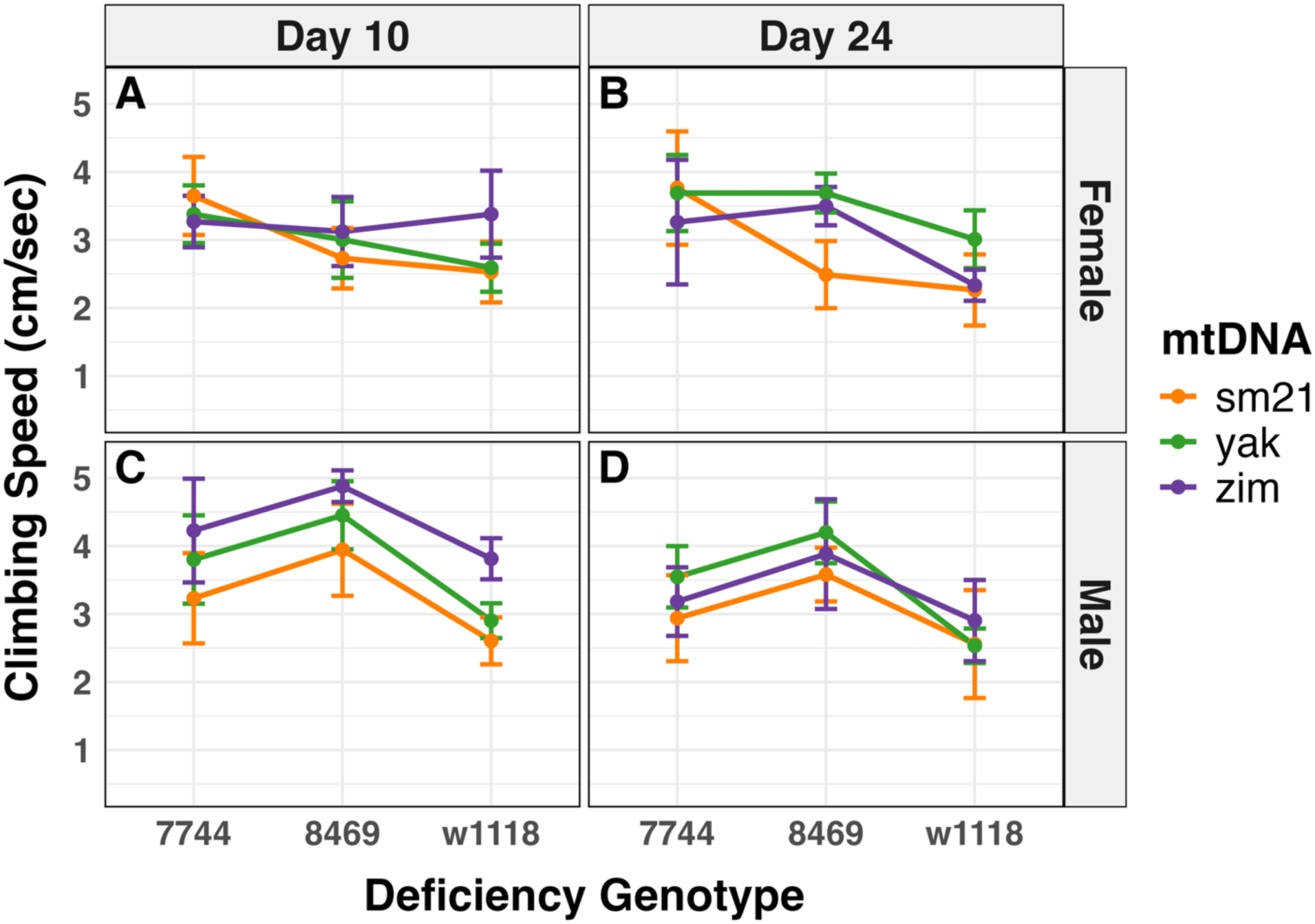
Climbing speed in block 3. Females show significant mtDNA, nuclear and mitonuclear effects, while male mitonuclear effects are not significant. Male mtDNA variation is greater than that in females.

**Tables 4A,B.** Analysis of variation for climbing speed in block 3 among 9 mitonuclear genotypes of females and males.

| FEMALES | Df | Sum Sq | Mean Sq | F value | Pr(>F) |  |
| --- | --- | --- | --- | --- | --- | --- |
| mito | 2 | 2.031 | 1.015 | 4.095 | 0.01986 | * |
| geno | 2 | 12.025 | 6.012 | 24.246 | 3.77e-09 | *** |
| day | 1 | 0.039 | 0.039 | 0.159 | 0.69078 |  |
| mito:geno | 4 | 4.789 | 1.197 | 4.829 | 0.00142 | ** |
| mito:day | 2 | 2.575 | 1.288 | 5.193 | 0.00734 | ** |
| geno:day | 2 | 1.603 | 0.802 | 3.232 | 0.04408 | * |
| mito:geno:day | 4 | 2.130 | 0.532 | 2.147 | 0.08144 | . |
| Residuals | 90 | 22.318 | 0.248 |  |  |  |

| MALES | Df | Sum Sq | Mean Sq | F value | Pr(>F) |  |
| --- | --- | --- | --- | --- | --- | --- |
| mito | 2 | 8.331 | 4.165 | 15.280 | 1.94e-06 | *** |
| geno | 2 | 29.079 | 14.539 | 53.337 | 5.28e-16 | *** |
| day | 1 | 6.845 | 6.845 | 25.110 | 2.69e-06 | *** |
| mito:geno | 4 | 1.540 | 0.385 | 1.412 | 0.23643 |  |
| mito:day | 2 | 3.129 | 1.564 | 5.738 | 0.00451 | ** |
| geno:day | 2 | 0.052 | 0.026 | 0.095 | 0.90922 |  |
| mito:geno:day | 4 | 0.169 | 0.042 | 0.155 | 0.96016 |  |
| Residuals | 90 | 24.534 | 0.273 |  |  |  |

Signif. codes: 0 '\*\*\*' 0.001 '\*\*' 0.01 '.' 0.1 ' ' 1
'mito' = mtDNA genotype, 'geno' = nuclear deficiency genotype, 'day' = age

The test of the MC hypothesis for climbing speed in block 3 is shown in Figure 3D. There are three nuclear backgrounds and two ages resulting in six CVmt pairs of points. This block shows more points above the 1:1 line, a result that is consistent with the MC hypothesis. Bootstrap resampling analyses (supplementary Figure S6) support the pattern seen in the MC test biplot (Figure 3D). Across the three experimental blocks for climbing speed there is no consistent pattern that the CVmt for males is greater than the CVmt for females, a pattern that does not support the MC hypothesis (Figures 3A-D). Some blocks show a trend towards a strong rejection of the MC hypothesis (Block 1, Figure 3A,B), while other blocks show a trend towards support of the MC hypothesis (Block 3, Figure 3D).

A general pattern that is apparent across these three blocks of climbing analyses is that the divergent mtDNAs from *D. simulans* and *D. yakuba* do not generate consistent negative effects in most of the nuclear backgrounds. The *siI, sm21,* or *yak* mtDNAs are frequently nested among the other mtDNAs for average climbing speed values. The placement of these outgroup mtDNAs is variable across nuclear backgrounds and experimental blocks: in some cases, the *yak* mtDNA is among the faster-climbing mtDNAs while the *sm21* mtDNA can be the slowest (Figure 5B,D). Given the ∼100-170 amino acid substitutions, and ∼600-960 total nucleotide substitutions between these outgroups and *D. melanogaster* mtDNAs (*Zim53* or *w1118*, see Figure 1 and Table 1), the degree of molecular divergence does not make these mtDNA genotypes ‘outgroups’ at a phenotypic level.

### Flight performance

The flight assay was performed on four nuclear backgrounds: three different heterozygous deficiency backgrounds (1491, 4959, 7837) plus the reference *w^1118^* homozygous background (Figure 6A,B; Table 5). The mtDNA effects on flight performance are significant in females but not in males. The nuclear background effect is strongly significant in both females and males, and the mitonuclear interaction term is significant in both sexes. The F-values for the mtDNA and mitonuclear interactions are larger in females than in males (Table 5). The mitonuclear genotype of *w^1118^* mtDNA on its native *w^1118^* nuclear background did not produce sufficient adults for a robust flight assay. We repeated the statistical analyses with *w^1118^* mtDNA removed from all nuclear backgrounds (a 3 mtDNA x 4 nuclear background analysis) and obtained qualitatively similar results with slight differences in the values of the ANOVA tables. See Tables 4A,B vs. 4C,D and Supplementary Figures S7 and S8A, B, C. Reversing the order of the mtDNA and nuclear main effects in the model reduced the significance of mtDNA in females and increased that of mtDNA in males such that they were equivalent. However, a linear model permitting type III sum of squares produced a result similar to the original model of landing height ∼ mtDNA × nuclear DNA, where the significance and F-values for mtDNA were greater in females than males (see supplementary Table S5A-D).

**Figure 6.**
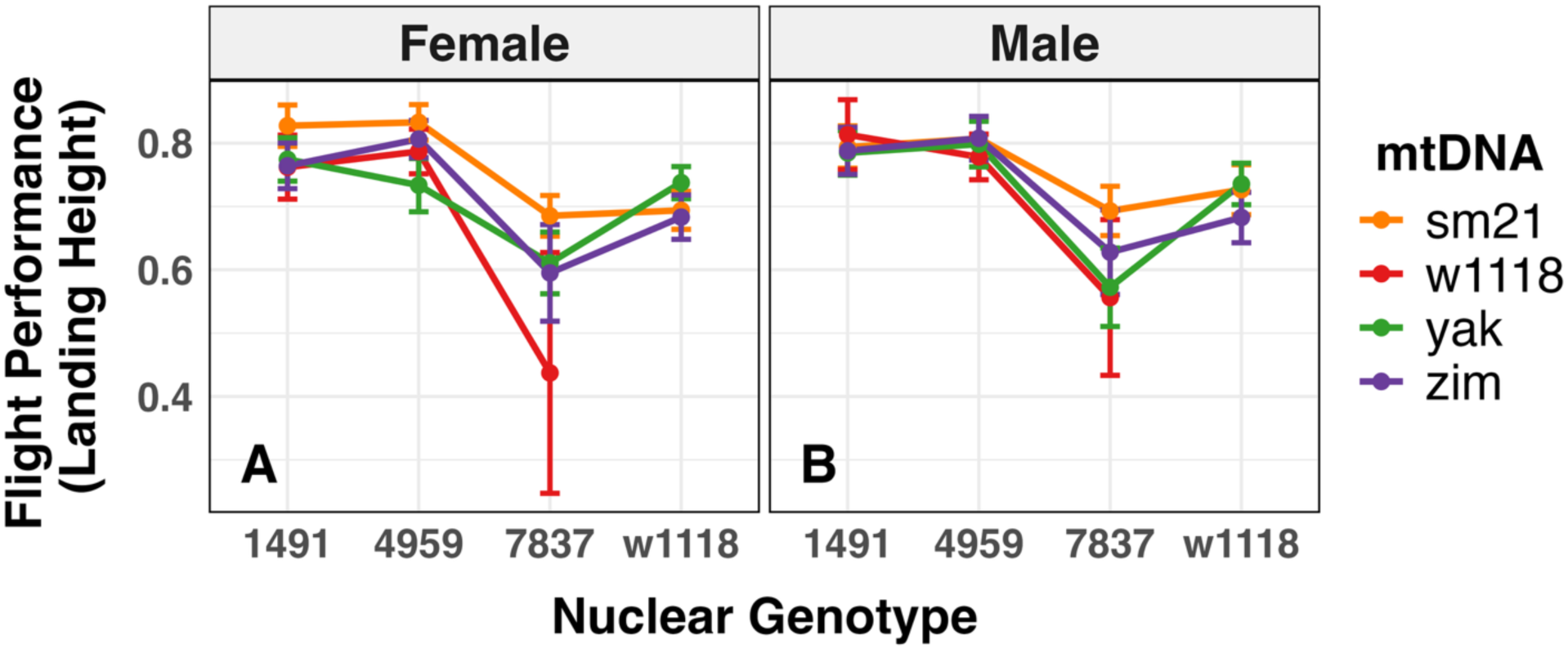
Flight performance of four mtDNAs across four nuclear backgrounds in females (A) and males (B).

**Table 5.** Analysis of variance for flight performance among females and males.

**A,B. 14 mitonuclear genotypes**
| FEMALES | Df | Sum Sq | Mean Sq | F value | Pr(>F) |  |
| --- | --- | --- | --- | --- | --- | --- |
| mito | 3 | 0.50 | 0.1677 | 4.179 | 0.00587 | ** |
| nuc | 3 | 6.36 | 2.1194 | 52.806 | < 2e-16 | *** |
| mito:nuc | 8 | 1.43 | 0.1783 | 4.442 | 2.44e-05 | *** |
| Residuals | 1630 | 65.42 | 0.0401 |  |  |  |

| MALES | Df | Sum Sq | Mean Sq | F value | Pr(>F) |  |
| --- | --- | --- | --- | --- | --- | --- |
| mito | 3 | 0.20 | 0.0659 | 1.454 | 0.22528 |  |
| nuc | 3 | 6.62 | 2.2068 | 48.674 | < 2e-16 | *** |
| mito:nuc | 8 | 0.93 | 0.1161 | 2.561 | 0.00894 | ** |
| Residuals | 1602 | 72.63 | 0.0453 |  |  |  |

**C,D. 12 mitonuclear genotypes**
| FEMALES | Df | Sum Sq | Mean Sq | F value | Pr(>F) |  |
| --- | --- | --- | --- | --- | --- | --- |
| mito | 2 | 0.45 | 0.2270 | 6.115 | 0.002269 | ** |
| nuc | 3 | 5.19 | 1.7296 | 46.602 | < 2e-16 | *** |
| mito:nuc | 6 | 0.96 | 0.1602 | 4.316 | 0.000254 | *** |
| Residuals | 1393 | 51.70 | 0.0371 |  |  |  |

| MALES | Df | Sum Sq | Mean Sq | F value | Pr(>F) |  |
| --- | --- | --- | --- | --- | --- | --- |
| mito | 2 | 0.13 | 0.0657 | 1.461 | 0.2325 |  |
| nuc | 3 | 5.59 | 1.8648 | 41.462 | <2e-16 | *** |
| mito:nuc | 6 | 0.73 | 0.1224 | 2.720 | 0.0125 | * |
| Residuals | 1397 | 62.83 | 0.0450 |  |  |  |

Df_7837 is the primary source of these effects, causing lower landing heights and wider variation across mtDNA in both males and females. While the nuclear backgrounds of these three deficiencies are not common across the non-deficient chromosomes, the hemizygosity in Df_7837 appears to have an epistatic effect across mtDNAs. It is notable that the Df_7837 was generated on a *w^1118^* stock background, and the mtDNA values for Df_7837 are very distinct from those on the homozygous *w^1118^* background in this block of flight assays, suggesting that the deficiency region plays an important role in the putative mitonuclear epistatic effect.

The biplot test of the MC hypothesis shows a slight trend towards more variation among females relative to males (Figure 3E). We note that the reduced data set alters the placement of one point in the biplot (Figure 3F), but this change does not alter the conclusion that the data are not consistent with the MC hypothesis.

As with climbing speed, the outgroup mtDNAs in *D. simulans* and *D. yakuba* do not cause strong defects in the flight performance. The biplot of CVmt for females and males do not support the MC hypothesis and though there are only four nuclear backgrounds to compare, the data more strongly support rejecting the MC hypothesis.

### Starvation resistance

Resistance to starvation was quantified for four mtDNAs on each of two different heterozygous deficiency backgrounds (7744 and 8469) plus the reference *w^1118^* homozygous background in both females and males (Figure 7 shows survival curves and interaction plots). A fifth mtDNA (*D. simulans siI*) was assayed on just the heterozygous deficiency background 8469 but was not included in the statistical analyses. There were significant differences in starvation survival among mtDNAs on all three nuclear backgrounds with some pairs of mtDNA showing differences in one background but not in another. This was true for females but not males. For females, the nuclear backgrounds show striking differences in starvation, with the 8469 nuclear background exhibiting dramatically higher resistance than 7744 or the reference *w^1118^* homozygous background. Overall, mtDNA, nuclear background, and their interaction all had highly significant effects (Table 6A,B). Interestingly, the outgroup mtDNAs of *D. simulans* and *D. yakuba* showed higher starvation resistance on the two deficiency backgrounds but were intermediate between the two different *D. melanogaster* mtDNAs (OreR and Zim53) on these more ‘native’ *w^1118^* nuclear chromosome sets (Figure 7A-F).

**Figure 7.**
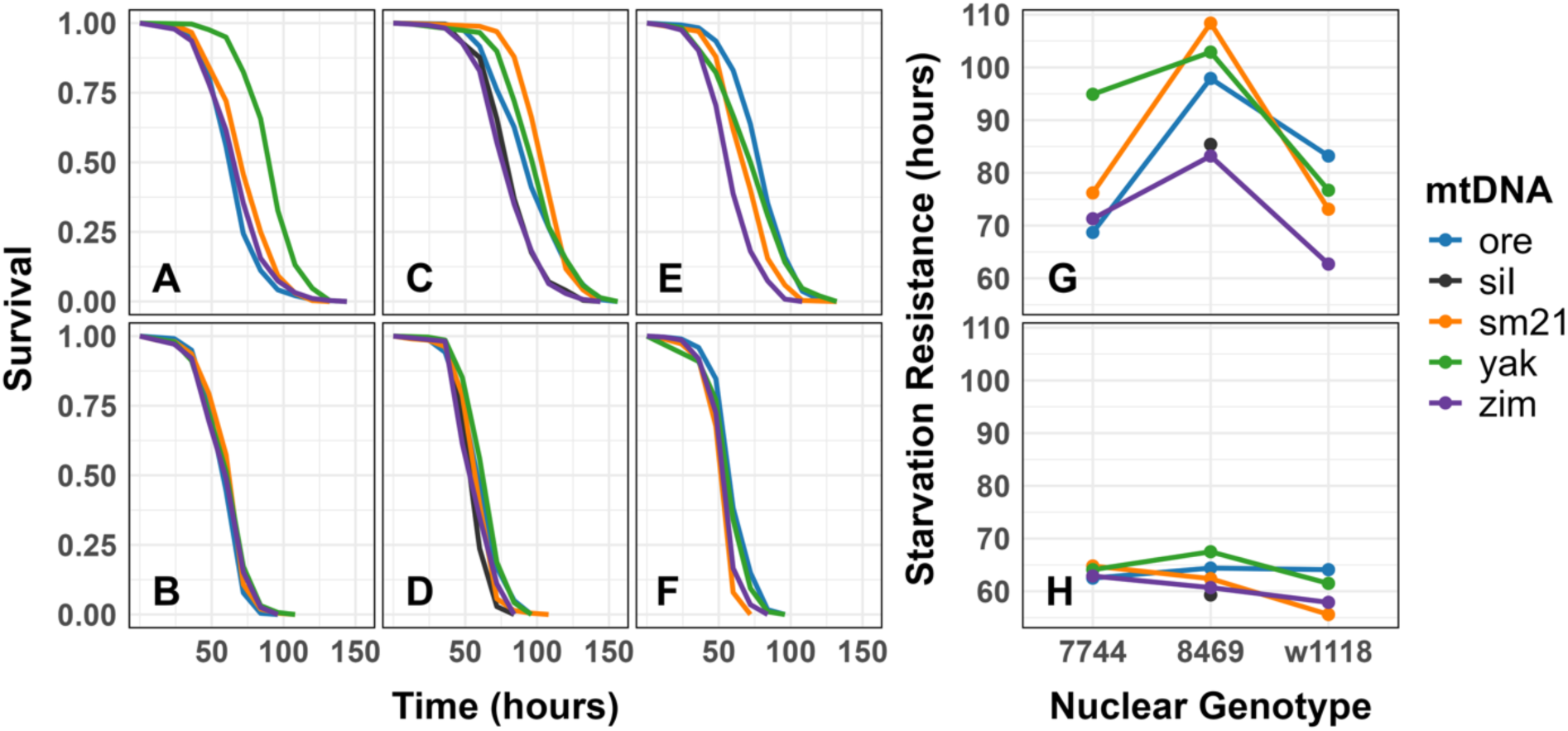
Survival curves and interaction plots for starvation resistance for female and male mitonuclear genotypes. A-F. Survival curves for four mtDNAs on three nuclear backgrounds (Females: A, C, E; males B, D, F). G and H. Interaction plots for mean starvation survival in females (G) and males (H).

**Table 6A,B.**
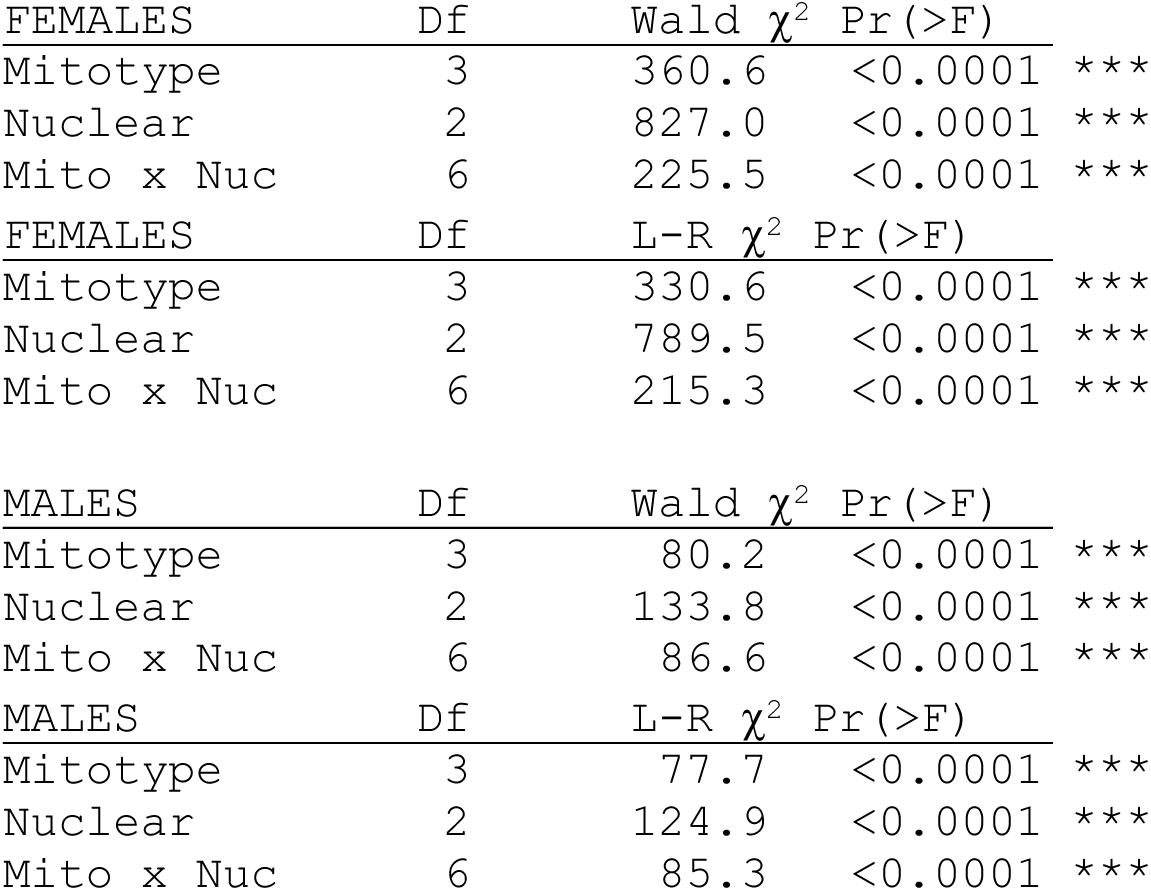
Parametric survival fits among 12 mitonuclear genotypes of females and males. The Wald test and a likelihood ratio test were determined separately for females and males, and the *siI* mtDNA was excluded as it was not paired with all three nuclear backgrounds.

Males showed strikingly lower starvation resistance and less variation across mtDNA or nuclear genetic backgrounds (Figure 7). As previously shown, males were much more sensitive to starvation than females usually attributed to the female egg mass that provides a source of nutrients that extends starvation resistance (Schwasinger-Schmidt et al. 2012). Despite reduced starvation resistance, there were significant effects of mtDNA, deficiency and their interaction in males, although the Ξ2 values were substantially lower than those of females (Table 5A,B, Figure 7). While there are only three different nuclear backgrounds in this assay, all three fall well below the 1:1 line in the MC biplot test, supporting rejection of the MC hypothesis (Figure 3G). Statistical support for the differences in male and female survival patterns in starvation is provided in supplementary Table S6.

### Test of the Mother’s Curse hypothesis

Taken together, the three MC test biplots from blocks 1-3 of the climbing assay and one each from flight and starvation assays show that the CVmt values for females are larger than those for males as more CV values fall below the 1:1 line than above it (Figure 8). While the difference in CV values between the sexes is not significant by a t-test (female mean = 0.130, s.d. 0.083, male mean = 0.099, s.d., 0.059, n=27) or a Wilcoxon signed rank test (W = 451, p-value = 0.1368), the slope of the line fit between these values does not include the 1:1 line in its confidence interval (Male-CVmt = 0.311×Female-CVmt + 0.058, F_1,25_ = 5.86, p < 0.0231; see Figure 8). These results are the opposite of what would be predicted under the MC hypothesis, indicating that the data not only fail to support it, but also falsify its primary prediction.

**Figure 8.**
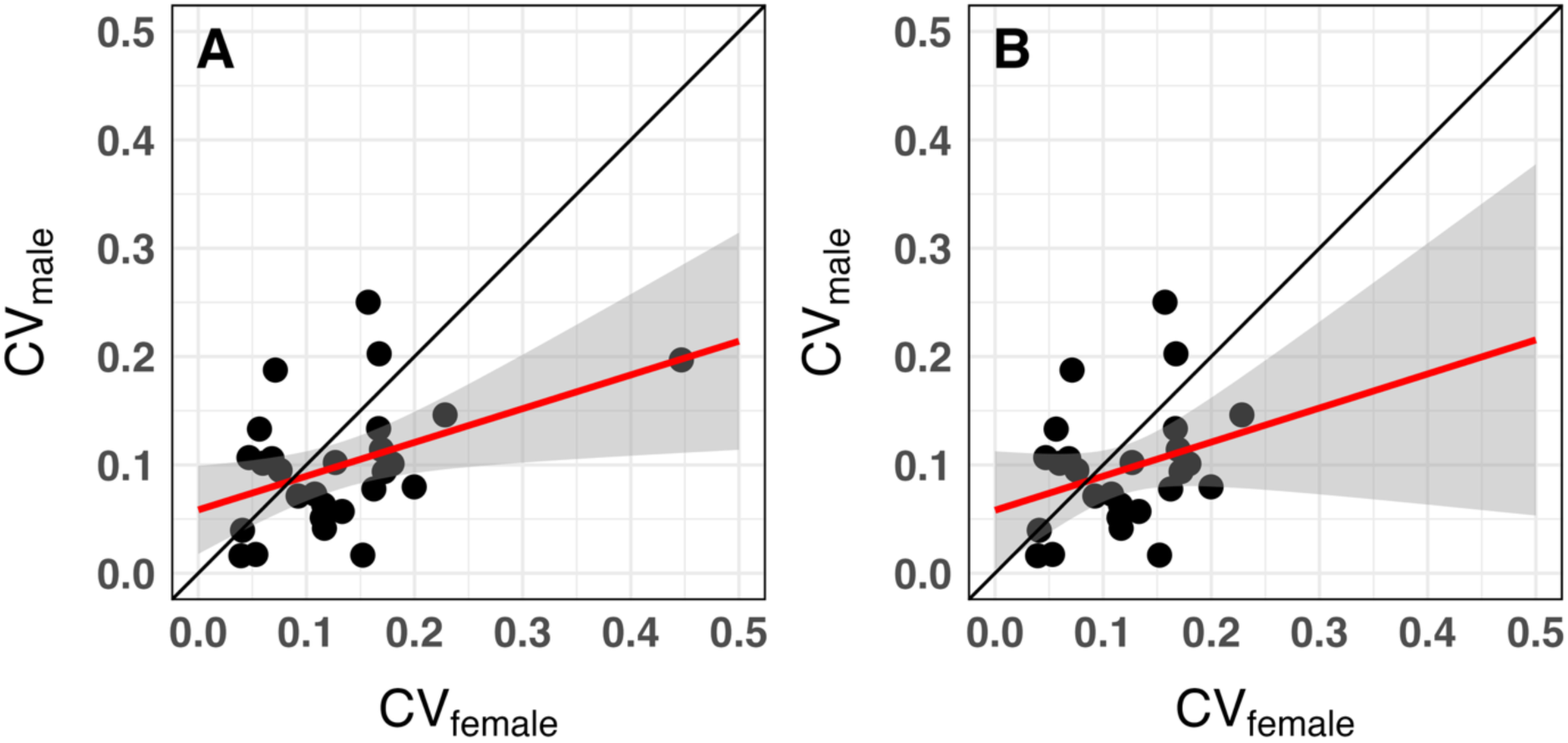
Test of the Mother’s Curse hypothesis across climbing, flight and starvation data. The coefficients of variation across mtDNA genotypes (CVmt) within each nuclear genotype or nuclear x age category for climbing, are plotted for females (x-axis, CV_female_) and males (y-axis, CV_male_). A. All CVmt values across the five data sets. B. Outlier point >0.4 for females removed. The red line is the best linear fit and the gray envelope defines the 95% confidence limits.

## Discussion

The maternal inheritance of mitochondrial genomes is consistently observed across the tree of eukaryotic life. This asymmetric mode of transmission suggests that mitochondria may have played a fundamental role in the origin of sex (Lane 2005), and mitochondria DNA (mtDNA) may be important source of sex-specific selection that continues in all populations (Camus et al. 2022). The latter topic is the focus of the Mother’s Curse concept, motivated by the work of (Cosmides and Tooby 1981, Frank and Hurst 1996) and popularized by (Gemmell et al. 2004). Since mtDNA is maternally inherited, natural selection can act on mtDNA-mediated phenotypic variation among females but cannot directly influence mtDNA effects in males. As a result, mtDNA mutations that are deleterious in males, but neutral or beneficial in females, can persist in populations. Accordingly, females should show lower levels of mtDNA-mediated phenotypic variation than males. Since mtDNA is a high-mutation-rate genome with no recombination, it is prone to Muller’s ratchet and mutational decay, which should result in reduced male fitness. The Mother’s Curse hypothesis (hereafter MCH) has been tested in a growing number of studies, some providing support (Rand et al. 2001, Sackton et al. 2003, Innocenti et al. 2011, Camus et al. 2012, Dowling and Adrian 2019, Carnegie et al. 2021), others showing mixed (Mossman et al. 2016, Montooth et al. 2019, Edmands 2024, Garlovsky et al. 2025), and even opposing results (Mossman et al. 2016, Mossman et al. 2017, Cayuela et al. 2023). The topic is more than an academic puzzle as the validity of the MCH could influence medical procedures such as mitochondrial replacement therapy or other treatments for mitochondrial disease (Reinhardt et al. 2013, Morrow et al. 2015, Eyre-Walker 2017, Hyslop et al. 2025, McFarland et al. 2025). The MCH has been invoked as a reliable genetic force in the proposal to eliminate disease vectors by driving extinction due to male decay (the ‘Trojan Female’ effect (Gemmell et al. 2013, Dowling et al. 2015)).

The current study seeks to address the generality of the MC process and to examine two issues that are potential limitations to empirical tests of the MCH that have been reported. Most studies use nuclear genomic backcrossing or chromosome replacement to place different mtDNAs onto one or more common, homozygous nuclear backgrounds. This provides a simplified nuclear context for the expression of mtDNA effects, but most sexually reproducing animals have measurable heterozygosity such that each individual has a unique nuclear genome. Even those studies that have used more than one nuclear background, the homozygous nature of the test may not be relevant to natural populations, including humans. Here we have addressed this concern using F1 mitonuclear genotypes that place different mtDNAs onto common heterozygous backgrounds carrying various deficiency chromosomes paired with the nuclear chromosomes from a standard *w^1118^* stock . Our approach assesses how different mtDNAs interact with nuclear genotypes that vary in the effects of gene dosage and additive and dominance factors.

A second issue concerns the long-term evolutionary consequence of MC. A strong, or adaptive, interpretation of the MCH posits that mtDNA mutations that are beneficial in females but deleterious in males should sweep through populations, causing persistent male decline. If this is a common phenomenon, one might be able to detect more pronounced effects of MC using mtDNAs that span a range of low, medium, and high levels of molecular evolutionary divergence. The absence of such effects would not support the adaptive scenarios of the MCH. We have conducted such a test of the MCH using mtDNAs from three different Drosophila species whose divergence spans <0.1 - ∼10 million years (*D. melanogaster, D. simulans* and *D. yakuba*). While the mtDNAs from the latter two species are not part of the breeding pool of *D. melanogaster*, we want to determine if the predictions of the ‘adaptive’ version of the MCH can be falsified. We have chosen three metabolically demanding performance traits that should challenge mitochondrial functions: climbing speed, flight performance and starvation resistance. All three traits are fitness-related and sexually dimorphic, but not sex-limited, so comparisons between sexes can provide a fair test of the MCH.

### Climbing speed

All three blocks of mitonuclear genotypes show significant effects of mtDNA and nuclear genotype in both males and females. In general, the impact of different nuclear genotypes greatly outweighs the effects of different mtDNA, as evident from F-values in Tables 2-6. This is to be expected based on vastly more nucleotides in a nuclear vs. an mtDNA genome. There are clear differences between males and females in the magnitudes of the effects of mtDNAs and in the presence of significant mito × nuclear (mitonuclear) interactions. With a couple of exceptions (e.g., block 3, Table 4), the effects of mtDNA among females are greater than that of males, and this trend is also apparent for the magnitude of mitonuclear epistatic interactions.

These results are not consistent with the simple prediction of the MCH. Admittedly the sample sizes of different mtDNAs and nuclear backgrounds are small within each experiment, but the replication of the vials and climbing events, in addition to the numbers of animals within each assay are substantial. While individual pairs of F-tests for the variance of female vs. male trait values are not significant, collectively the results indicate that despite significant genetic effects of mtDNAs, the signal is not clearly skewed in the direction of more phenotypic variation among mtDNA in males than females (Figure 3A-D).

Previous studies of climbing speed in Drosophila have identified significant mtDNA and mitonuclear effects (Sujkowski et al. 2019, Spierer et al. 2021). The climbing speed assay challenges leg muscles and some component of the startle response that drives negative geotaxis. The *tko* gene is particularly interesting: the defining mutation is in a nuclear encoded mitochondrial ribosomal protein and shows delayed recovery from a knock down assay using a laboratory vortex (Toivonen et al. 2001). This may impact mitochondrial functions associated with neuromuscular firing that allows flies to initiate their natural tendency to exhibit negative geotaxis. The climbing assay we use (FreeClimber; (Spierer et al. 2021)) has three gentle ‘knock-downs’ of ∼20 flies in each of six glass vial experiencing an 8 cm drop. Few, if any, flies are immobilized from the drop, but some combination of mitochondrial function in neuromuscular activity may contribute to the speed of climbing.

### Flight performance

The flight performance results show that nuclear genetic variation is the primary factor in this assay. While females show significant mtDNA effects, males show no significant mtDNA effects in both treatments of the data sets. However, males and females both showed significant mitonuclear interactions. The results show a trend of higher CVmt values in females, but the sample of tests is small, precluding a meaningful F-test.

The previous studies of flight performance in Drosophila identified a number of mitonuclear statistical interactions (Sujkowski et al. 2019), and effects of neurodevelopmental genes (Spierer et al. 2021) that contribute to performance, but the latter study did not identify significant mitochondrial signals in the GWAS analyses. This particular flight assay is an acute response to an abrupt drop into the flight column and may challenge more rapid neuromuscular response functions than continuous mitochondrial respiration requiring proper mitonuclear genetic interactions. Flight is an important fitness trait that requires metabolically active flight muscle rich in mitochondria, so it should be a viable candidate for a test of the MCH.

### Starvation resistance

The starvation resistance data provide a clear falsification of the predictions of the MCH. MtDNA genetic variation has a substantial and significant effect in females, but a very limited one in males. Starvation resistance challenges the anabolic and catabolic functions of mitochondria: the flies are provisioned on normal food and allowed to accumulate metabolic storage products. When assayed, long lived genotypes are those that have larger reserves or metabolize their reserves at a lower rate. These functions certainly include pathways that run through glycolysis, the TCA cycle, OXPHS, beta-oxidation, among others that require proper mitochondrial function, and presumably proper mitonuclear coordination of gene products.

### Mitonuclear interactions with chromosomal heterozygosity and deficiencies

A primary goal of this study was to determine if the phenotypic effects of alternative mtDNAs could rise above the phenotypic effects of different nuclear backgrounds heterozygous for chromosomal deletions examined in each assay. It may be that the nuclear backgrounds we chose to employ – replicable heterozygotes with a chromosomal deficiency in chromosome 2 – are not good proxies for natural nuclear polymorphism. However, structural variation, gene dosage, and the additive and dominance effects of heterozygosity are all important components of quantitative genetic variation in natural populations, and the current model seeks to explore that complex space. The deficiencies differed in size and thus lacked variable numbers of genes (23-146 loci; see Table S1). They were chosen as a sample of chromosome 2L which harbors a typical density of OXPHOS genes. While some deficiencies have loci important in mitochondrial functions (e.g., mEFTs, mitochondrial translation elongation factor Ts in Df 7837; mTTF, mitochondrial transcription termination factor in Df 9061), there are many uncharacterized genes that may have little direct interaction with mtDNA encoded genes. Nevertheless, the results showed that there is clear evidence for mitonuclear epistatic interactions between the mtDNAs studied and the different Df backgrounds. This further underscores the challenge of understanding the significance of phenotypic effects of mtDNA variation in homozygous nuclear backgrounds.

### Generality of the Mother’s Curse phenomenon

Figure 8 provides a clear picture of the collective results across these different assays. The trend of this biplot test of the MCH is opposite in direction of the primary prediction: there is higher mtDNA-mediated phenotypic variation among females than males. As such the data not only fail to support the MCH but actually falsify it. While the difference between the CVmt values for males and female is not significant, the trend is in the wrong direction and does not include the null expectation of a 1:1 slope within the confidence limits. These results are in stark contrast to other studies that report clear signals of the MC pattern, in some cases using the same nuclear strain we have used as a standard nuclear background, *w^1118^* .

### MtDNA divergence and Mother’s Curse

The data repeatedly show that the divergence of mtDNAs from *D. simulans* and *D. yakuba* does not provide strong evidence for disrupted coadaptation between the nuclear genome of *D. melanogaster* and the mtDNAs from the other species that have not interacted with one another for millions of years. This implies some kind of stabilizing selection that also does not induce unruly mitonuclear epistatic interactions (e.g., the foreign mtDNAs do not have wildly different crossing reaction norms in the interaction plots).

The many accumulated base pair and amino acid substitutions in these mtDNAs may have relatively mild impacts on the functioning of the OXPHOS complexes - the sites of putatively important contact between the gene products of these two genomes. Alternatively, there could have been compensatory mutations in nuclear genes that have accommodated deleterious mutations in mtDNA-encoded genes effectively rescuing or stabilizing the consequences of continuous mtDNA mutations (Rand et al. 2001, Rand et al. 2004, Adrion et al. 2016a, Hill 2020). Notably, there is no prominent signature of elevated dN/dS values or signatures of positive selection in the nuclear encoded OXPHOS genes across the phylogeny of the *D. melanogaster* species subgroup using codon tests in PAML (Drosophila 12 Genomes et al. 2007, Larracuente et al. 2008). Comparisons of mtDNA encoded proteins across the 12 Drosophila genomes reveals lower dN/dS values than for nuclear encoded proteins (Montooth et al. 2009). The higher dS values for mtDNA genes than nuclear genes are consistent with a history of purifying selection on mtDNA encoded proteins.

In previous studies, our group has failed to provide support for the MCH in much larger panels of mitonuclear genotypes (Mossman et al. 2016). These studies have been criticized for the use of limited number of intraspecific mtDNAs, three from *D. melanogaster* and three interspecific mtDNAs from *D. simulans*, arguing that the interspecific genotypes generate unnatural combination akin to a genotype being placed in a foreign environment (Dowling and Adrian 2019, Nagarajan-Radha et al. 2020). If this were true one would expect that these genotypes would accentuate the evidence for MC, or at least the significance of mtDNA based phenotypic effects. That these divergent mtDNAs are not phenotypic outliers in the performance traits we have assayed in this study indicates that there is, surprisingly, nothing unusual about these mtDNA haplotypes in terms of overall function. These results are not consistent with the adaptive version of the MCH: males carrying mtDNA that diverged ∼10 million years ago show no evidence of reduced performance (Montooth et al. 2010). By comparison, the evidence for the breakdown of mitonuclear coadaptation in *Tigriopus californicus* is compelling where the genetic divergence between intraspecific mtDNAs of *Tigriopus californicus* is as great as between orders of mammals (Ellison and Burton 2008, Burton 2022). Notably, not all pairs of reciprocal crosses that generate disrupted mitonuclear genotypes in that system show the same magnitude of effects (Healy and Burton 2020, Healy and Burton 2023). One hypothesis that may reconcile the different results from *Tigriopus* and *Drosophila* is the difference in effective population size resulting in stronger purifying selection in *Drosophila*. Deleterious mutations in *Tigriopus* are more likely to persist or fix, while they are likely to be purged more effectively in *Drosophila*. Recent studies further show that there is both positive (Li et al. 2024) and negative (Watson et al. 2022) evidence for the Mother’s Curse patterns in *Tigriopus* (Edmands 2024).

Interestingly, *Tigriopus* does not have distinct sex chromosomes and may provide a more ‘fair’ nuclear background for tests of MCH. The question remains whether the MCH patterns in *Drosophila* could stem from the hemizygosity of the X chromosome in males, which could expose more recessive phenotypic variation in males than in XX females, without any contribution from mtDNA. The enhanced co-transmission of mtDNA and X-linked variation provides strong opportunities for nuclear effects that covary with mtDNAs (Rand et al. 2001), so tests of the MCH with different sex chromosome systems may be informative.

### How should we test the Mother’s Curse hypothesis?

The Mother’s Curse is an entirely logical idea with a clear prediction and compelling evidence that it can work in some cases (e.g., cytoplasmic male sterility in plants). The fact that some data support the MCH and other data do not, or even falsify it, means that the MCH is not a consistent genetic effect, i.e., it can be idiosyncratic (Garlovsky et al. 2025). For example, Haldane’s rule holds in the vast majority of taxa, even in female heterogametic species, demonstrating that the genetic bases are strong and repeatable even if there is not one mechanism (Coyne 1985, Orr 1993, Delph and Demuth 2016, Cowell 2023). Garlovsky et al. (2025) provide a compelling test for mitonuclear interactions, coevolution, and Mother’s Curse using rigorous statistical analyses and show that mitonuclear effects among three female reproductive traits and nine male fertility traits are more apparent in males than in females, suggesting that males are more sensitive to mtDNA variation. The main limitation of this conclusion with respect to the MCH is that the significant effects are limited to male traits associated with sperm function - traits that are not present in females (even though sperm function can influence female reproduction). The difference between males and females in the effects of mtDNAs is central to the MCH so additional tests of this hypothesis should focus on traits that are not restricted to one sex. It would be interesting to know, among the genotypes tested by Garlovsky et al. (2025), if fitness traits shared by both males and females also show mtDNA sensitivity in females. The results of the current study support the notion that mitonuclear interactions and MC effects are idiosyncratic, but the overall conclusions clearly falsify the main prediction of the MCH. Mother’s Curse is a compelling idea, but it should remain a hypothesis to be tested against other sex-specific genetic causal factors that might curse the sex unable to transmit the mitochondrial (or organelle) genome.

## Data Availability

Drosophila strains, code and raw data are available at the Rand Lab GitHub site for this publication (https://github.com/DavidRandLab/Absence-of-Mothers-Curse). The authors affirm that all analyzed data necessary for confirming the conclusions of the article are present within the article, figures, tables and GitHub site.

## Author contributions

DMR conceived and designed the study, collected and analyzed data, wrote the manuscript. FAL, KMB, LM collected data. LJD and YR provided statistical analyses and comments on the manuscript.

## Acknowledgements

The authors thank Damian Dowling, Klaus Reinhardt, Justin Havird and Suzanne Edmands for comments on an earlier draft. Several attendees of PEQG 2022, DROS 2023 contributed feedback on poster versions of these results.

## Supplementary Figures

**Supplementary Figure S1.**
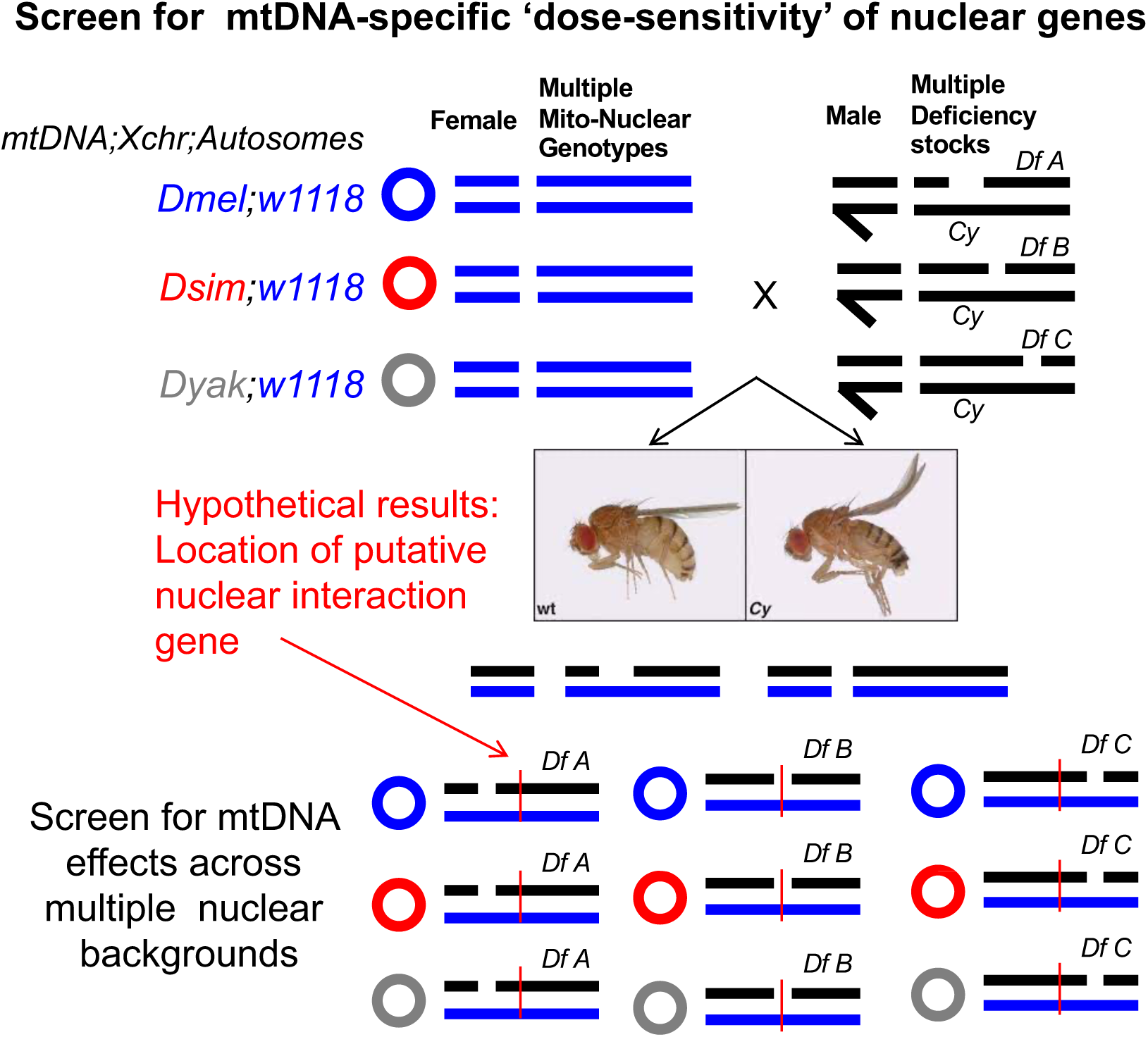
Crosses between female mitonuclear stocks and male deficiency (Df) stocks heterozygous for different deficient 2^nd^ chromosomes and a marked balancer chromosome carrying the curly wing marker (CyO). F1 offspring with wild type wing phenotypes are heterozygous for a common *w^1118^* 2^nd^ chromosome and different deficiency chromosomes. Comparisons across mtDNAs in common Df nuclear backgrounds and across Df backgrounds in common mtDNAs provide estimates of mitonuclear effects on phenotypes.

**Supplementary Figure S2.**
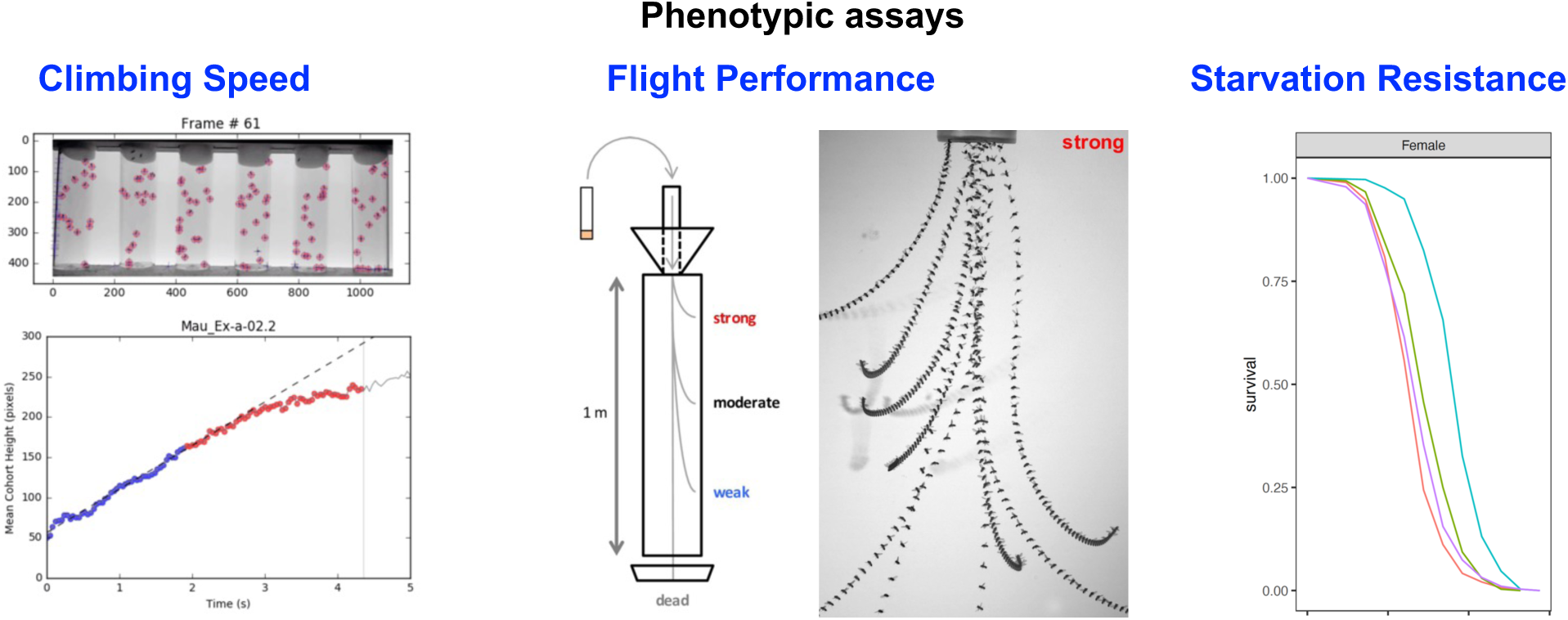
Examples of the three performance traits quantified for different mitonuclear genotypes paired with deficiency chromosomes (see Figure S1).

**Supplementary Figure S3.**
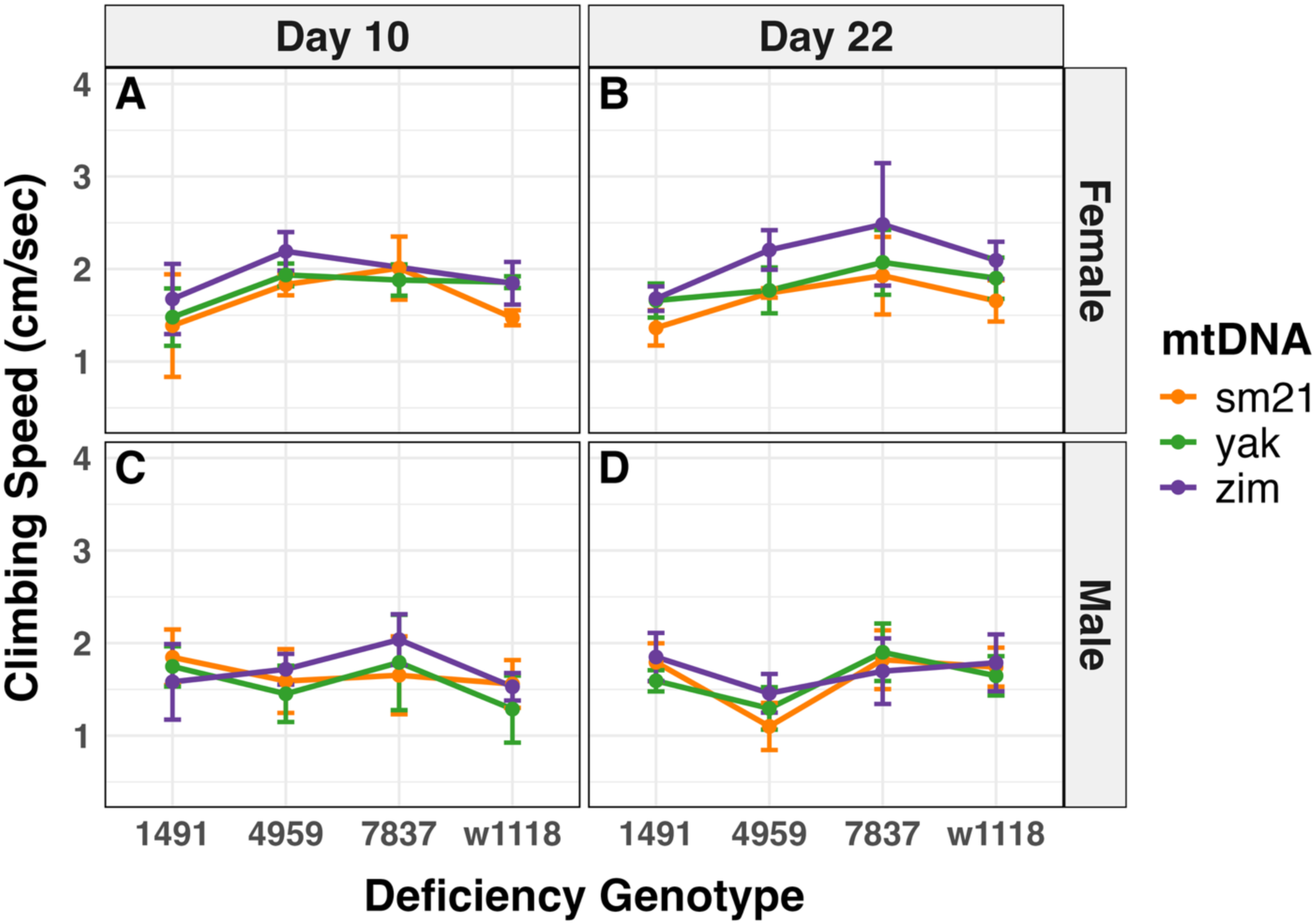
Climbing speed for block 1 using the reduced data set of 12 mitonuclear genotypes. Compare with Figure 2 from main text.

**Supplementary Figures S4A,B.**
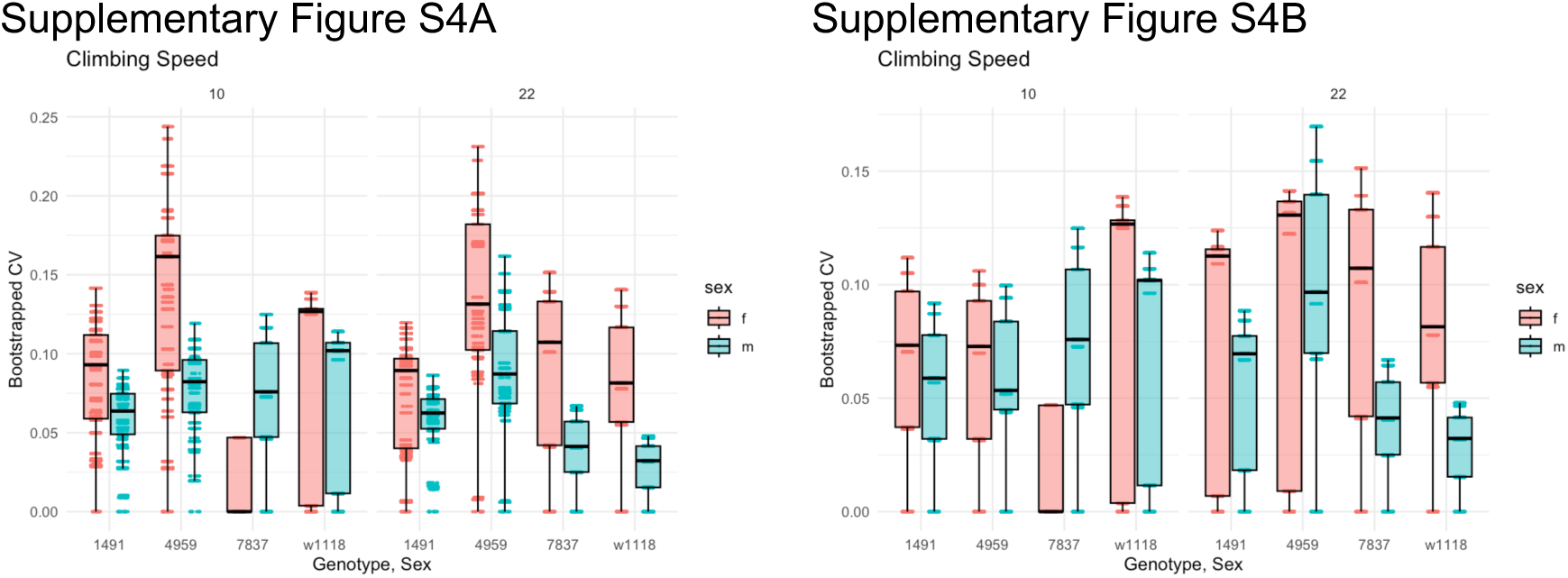
Bootstrap analyses of CVmt values from climbing speed block 1 full data set (S4A) and reduced data set (S4B). Resampling the mean climbing speeds confirm the patterns of CVmt variation between females (f) and males (m) in Figure 3A and B.

**Supplementary Figure S5.**
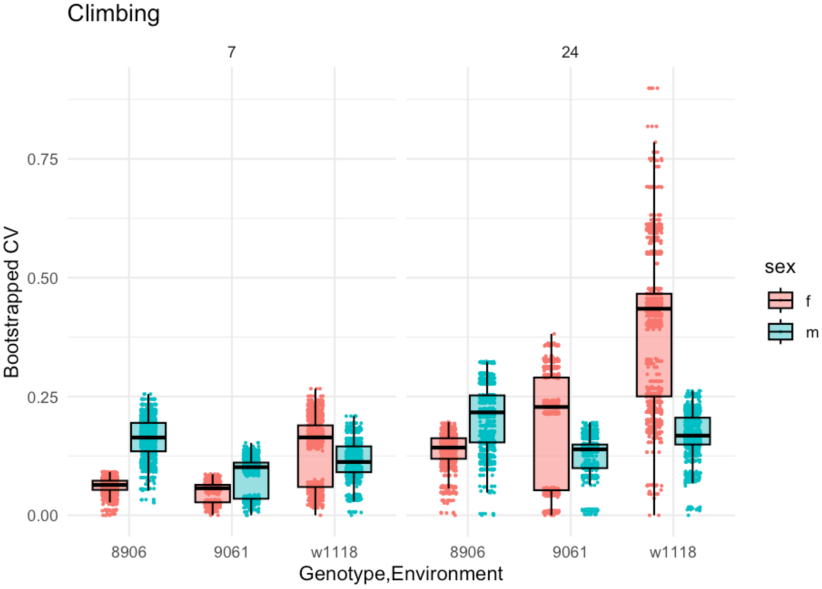
Bootstrap analysis of CVmt values for climbing speed, block 2. Resampling the mean climbing speeds confirm the patterns of CVmt variation between females (f) and males (m) shown in Figure 3C.

**Supplementary figure S6.**
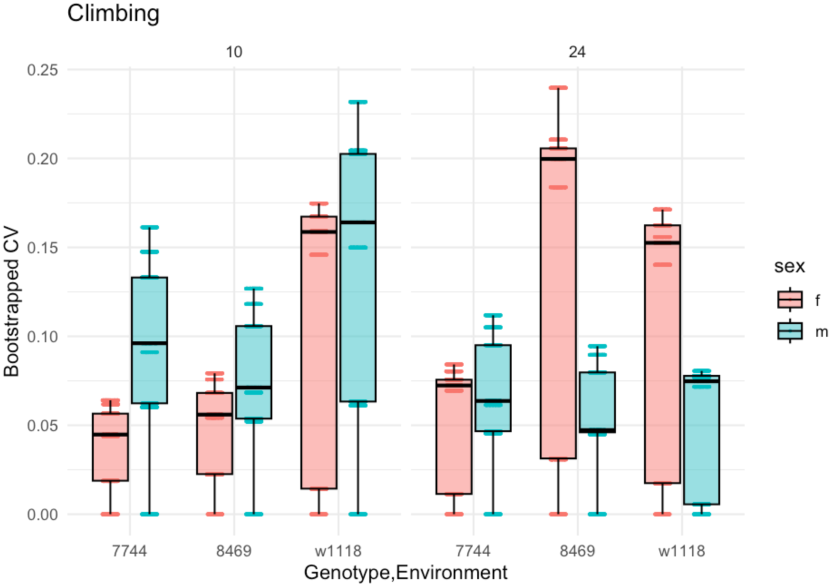
Bootstrap analysis of CVmt values for climbing speed, block 3. Resampling the mean climbing speeds confirm the patterns of CVmt variation between females (f) and males (m) shown in Figure 5.

**Supplementary figure S7A.**
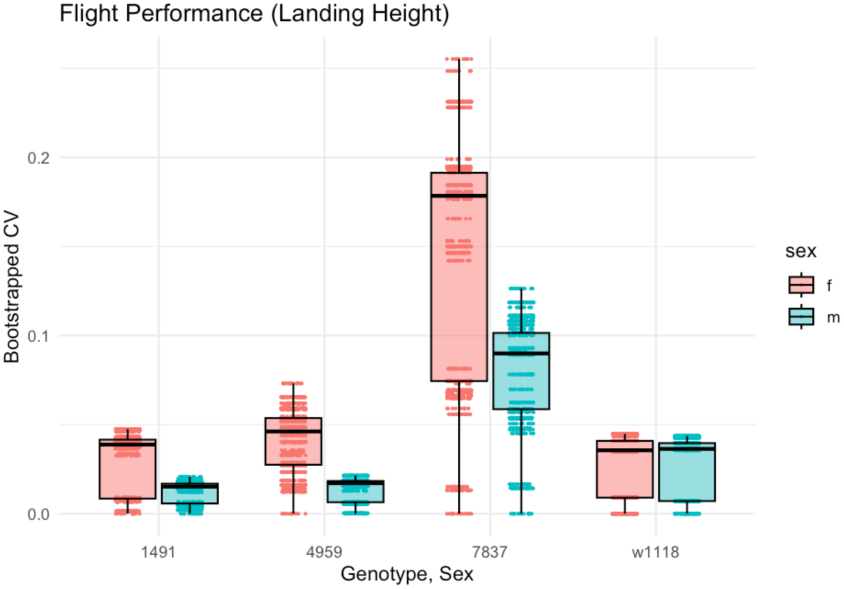
Bootstrap analysis of CVmt values for flight performance. Resampling the mean flight landing heights confirm the patterns of CVmt variation between females and males shown in Figure 6 and 3E.

**Supplementary figure S8A,B.**
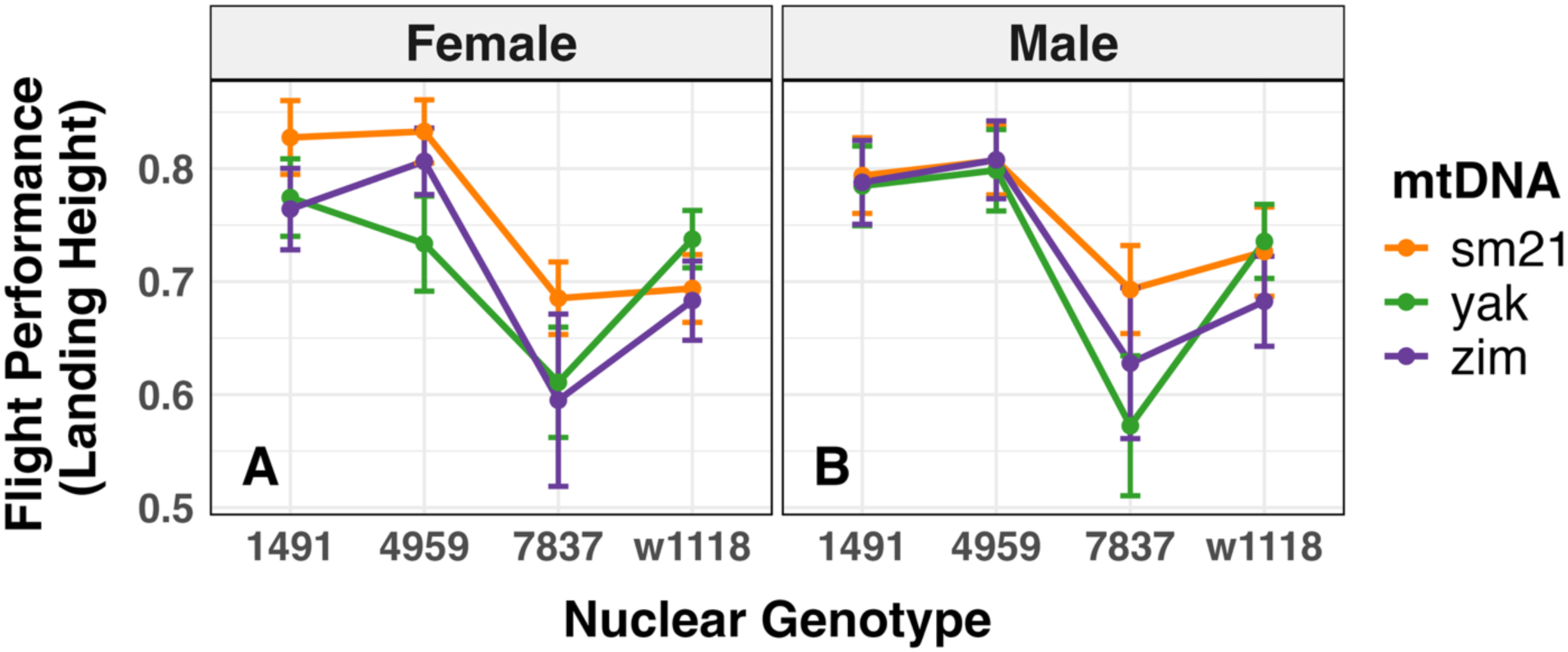
Flight performance results for the reduced data set of 3 mtDNA x 4 nuclear backgrounds. Interaction plots for females (S7B) and males (S7C). The biplot test of the MC hypothesis for this data set is shown in Figure 3F.

**Supplementary figure S8C.**
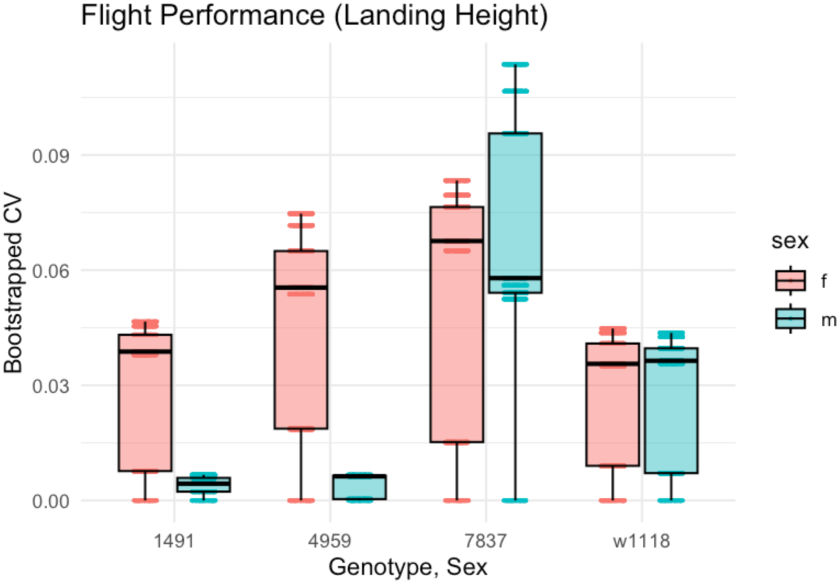
Bootstrap analysis of CVmt values for flight performance of reduced data set. Resampling the mean flight landing heights confirm the patterns of CVmt variation between females and males shown in Figure S8A, B.

**Supplementary Table S1.**
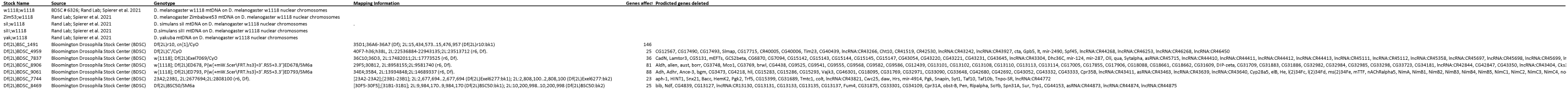
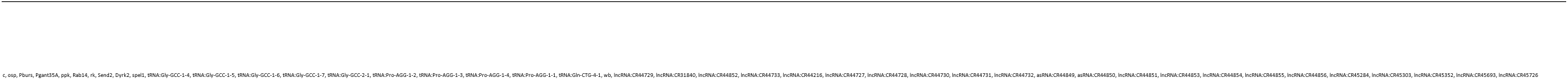

**Supplementary Table S2.**
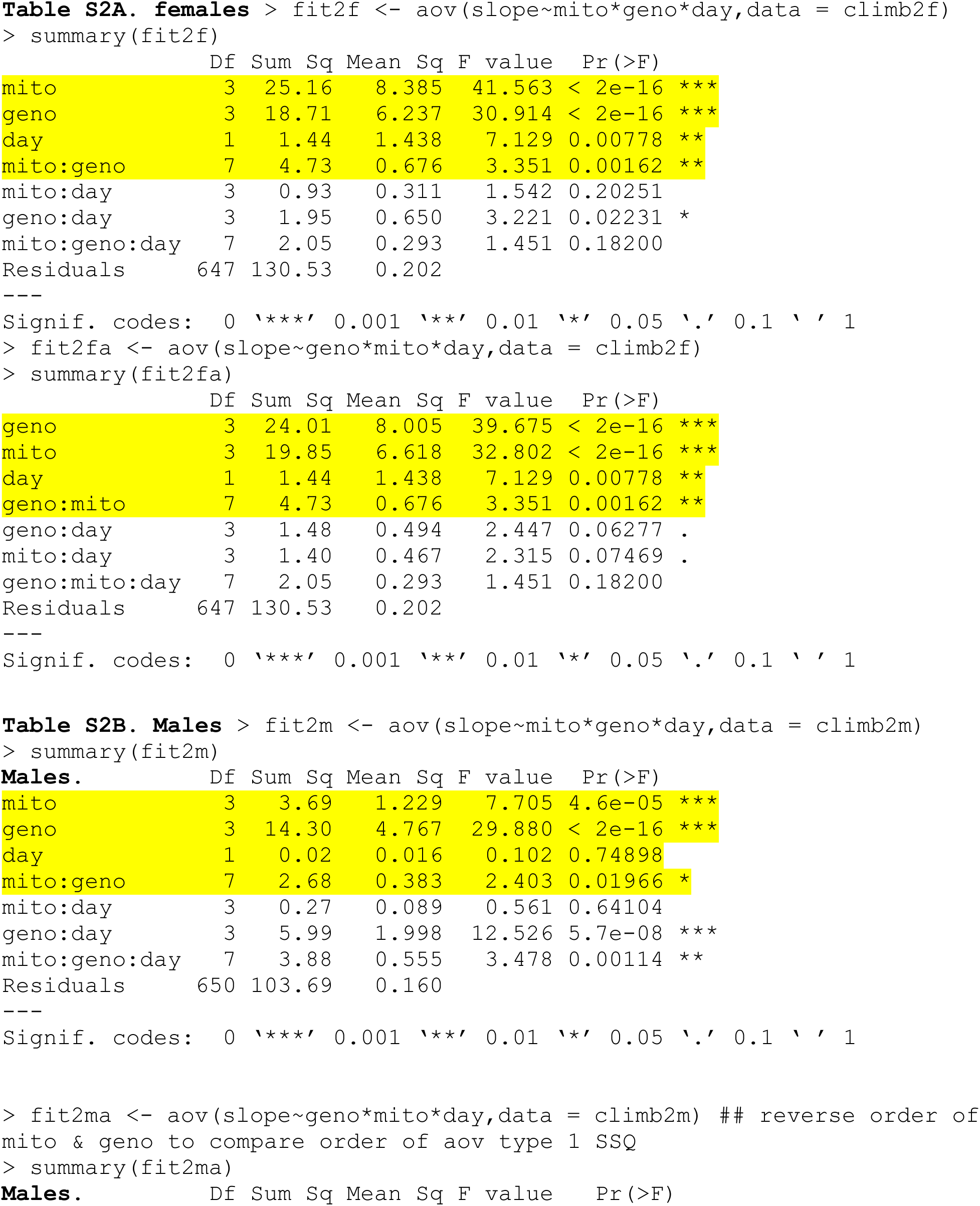
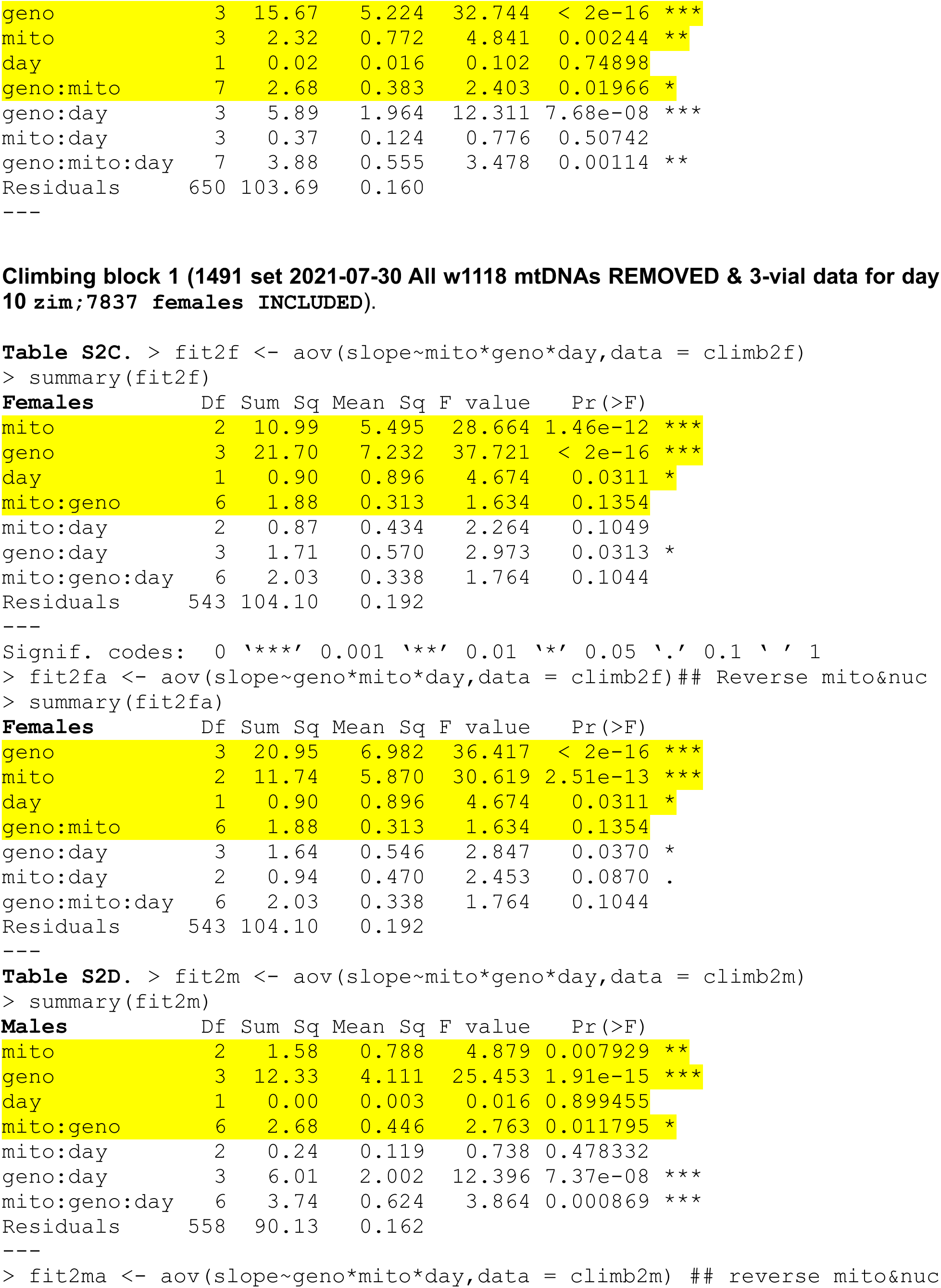
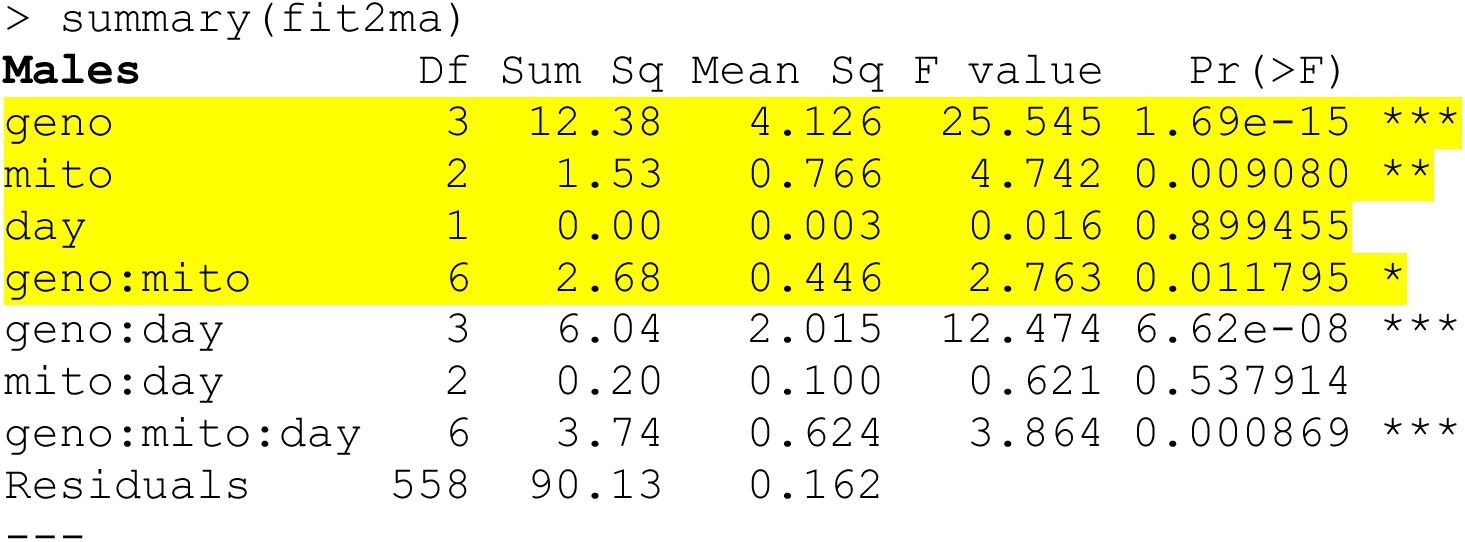
Climbing block 1 (1491 set 2021-07-30 with w1118 mtDNA present and 3-vial data for day 10 zim;7837 females INCLUDED)

**Supplementary Table S3.**
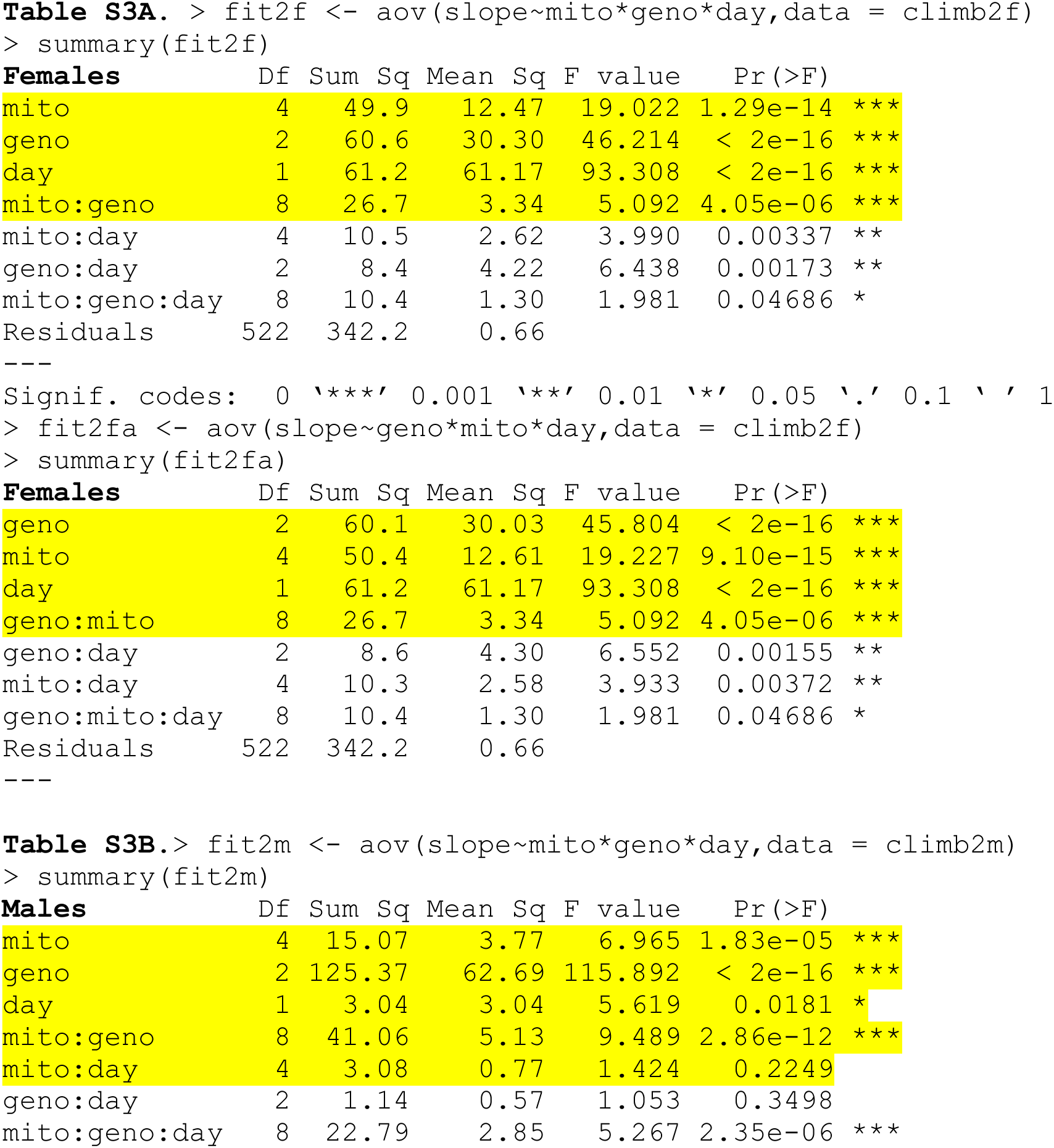
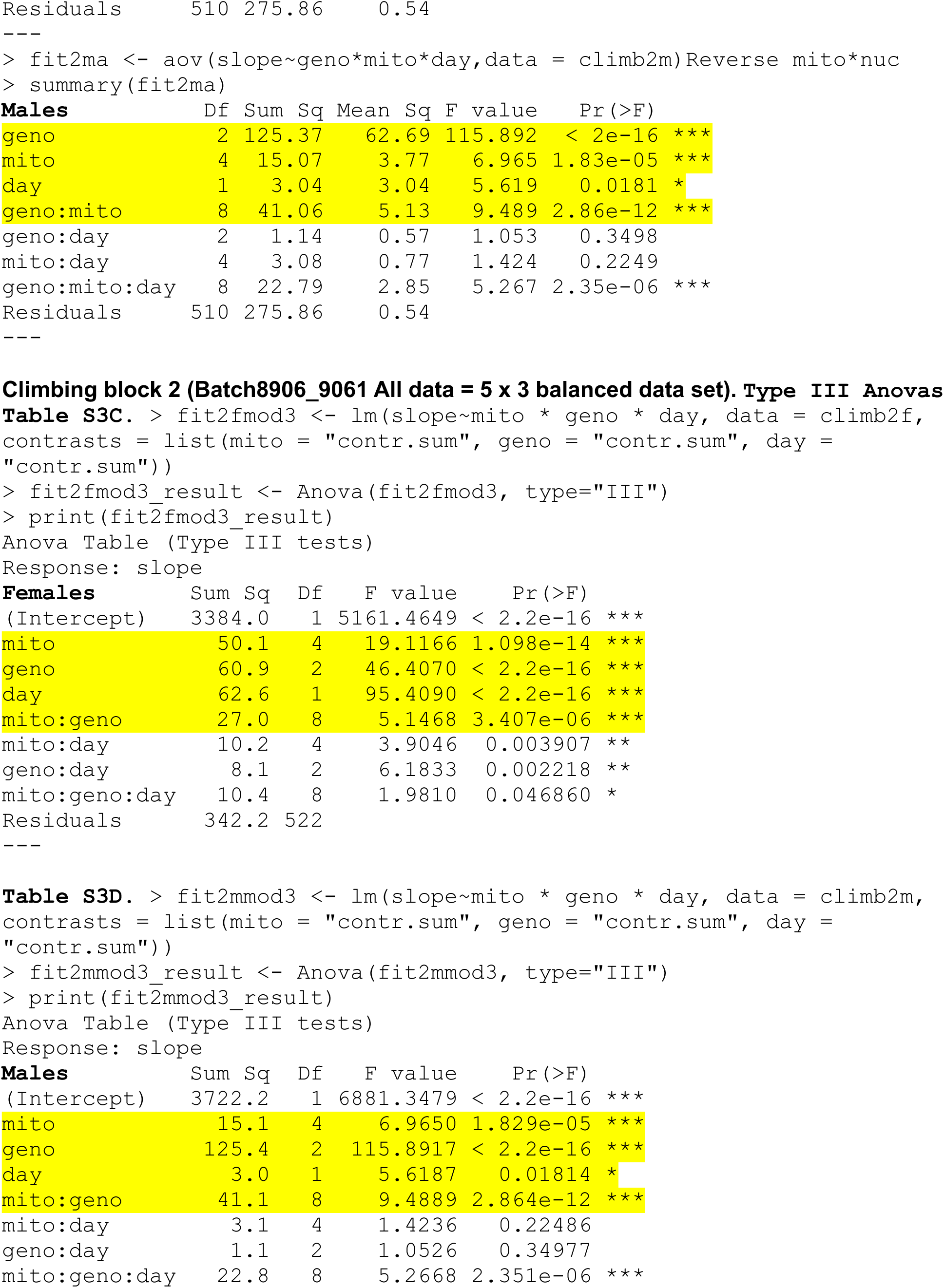
Climbing block 2 (Batch8906_9061 All data = 5 × 3 balanced data set).

**Supplementary Table S4.**
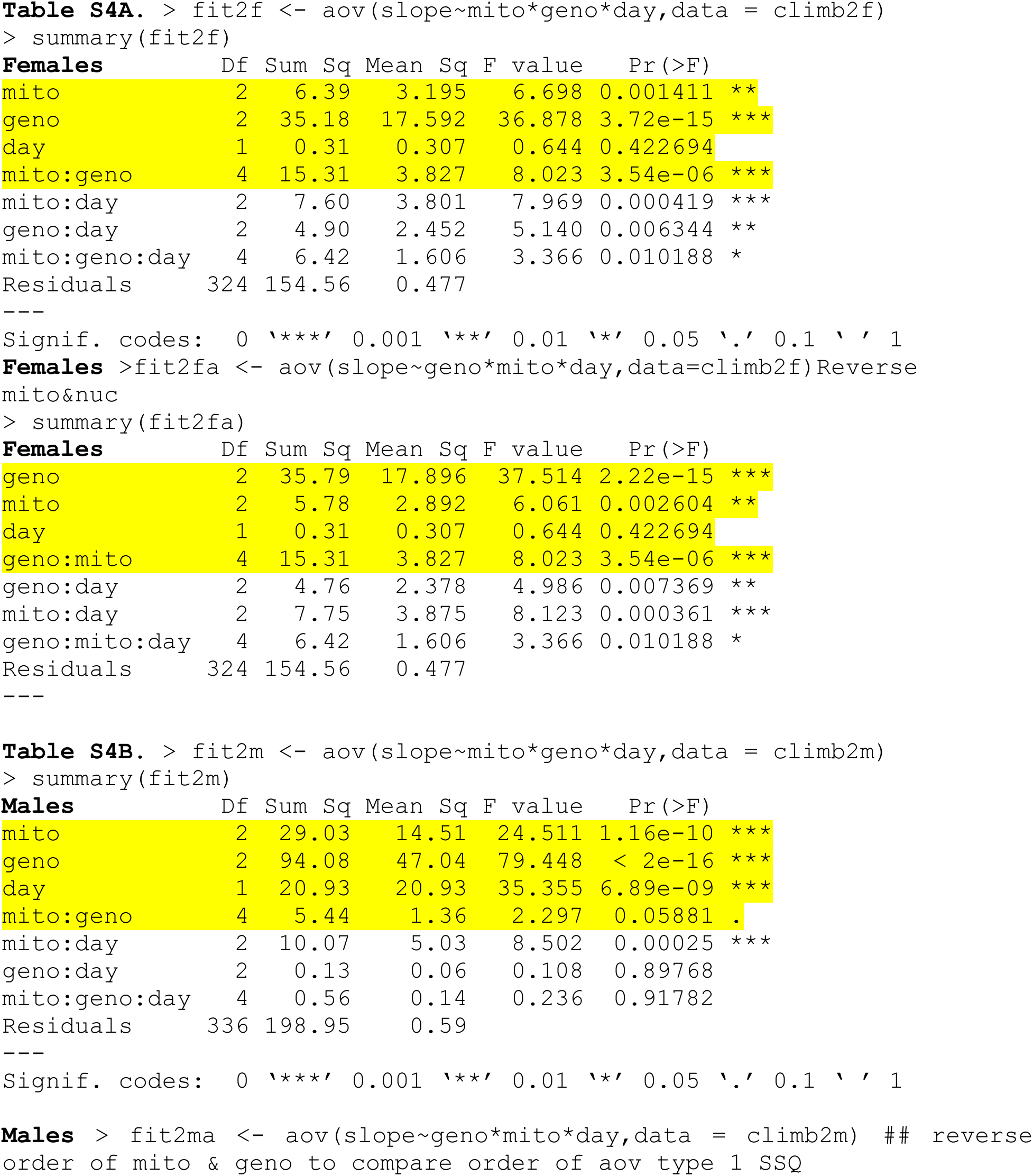
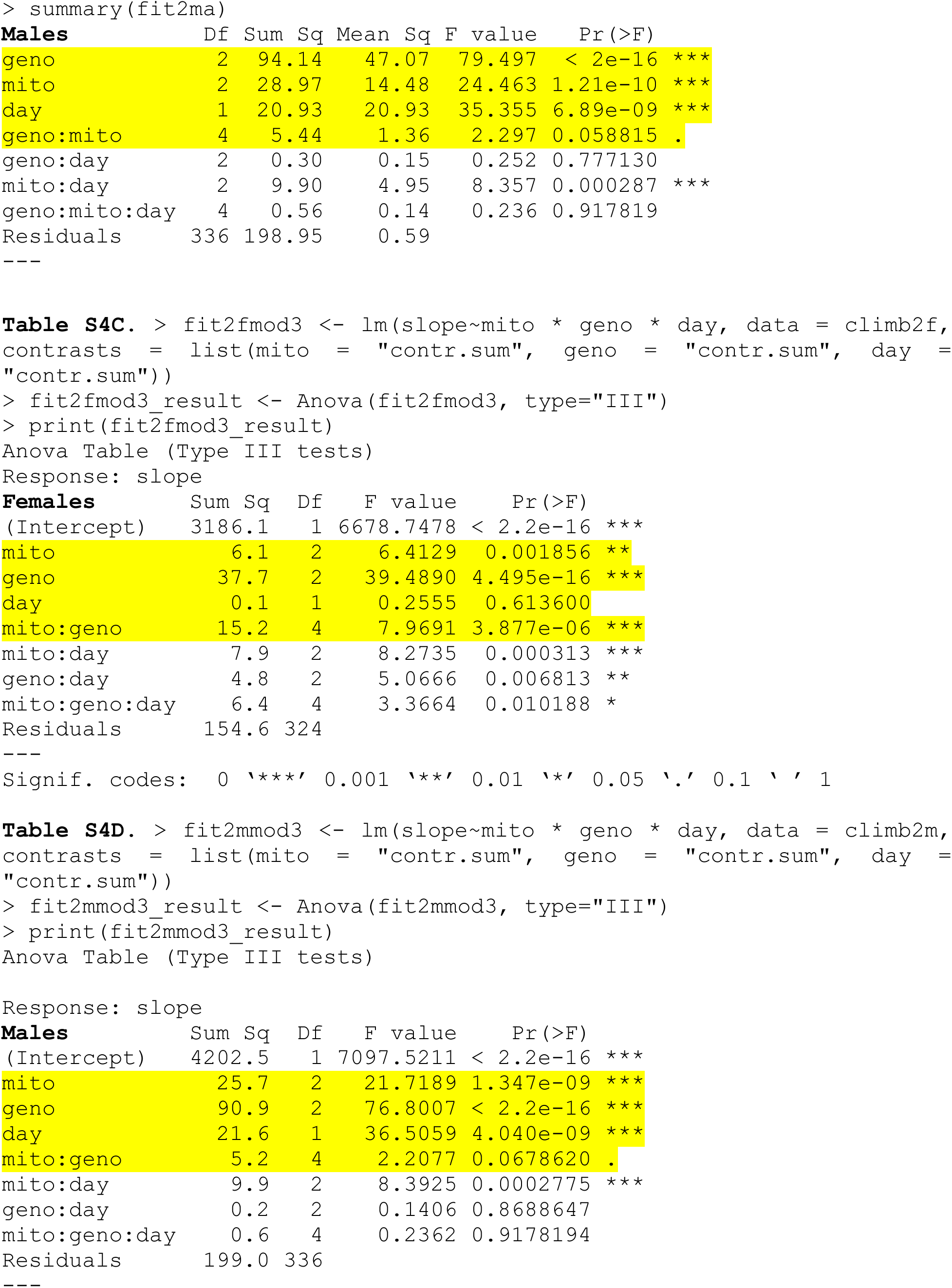
Climbing block3 (Data for 2021_07_15 batch of Dfs 7744 8469 w1118)

**Supplementary Tabl S5.**
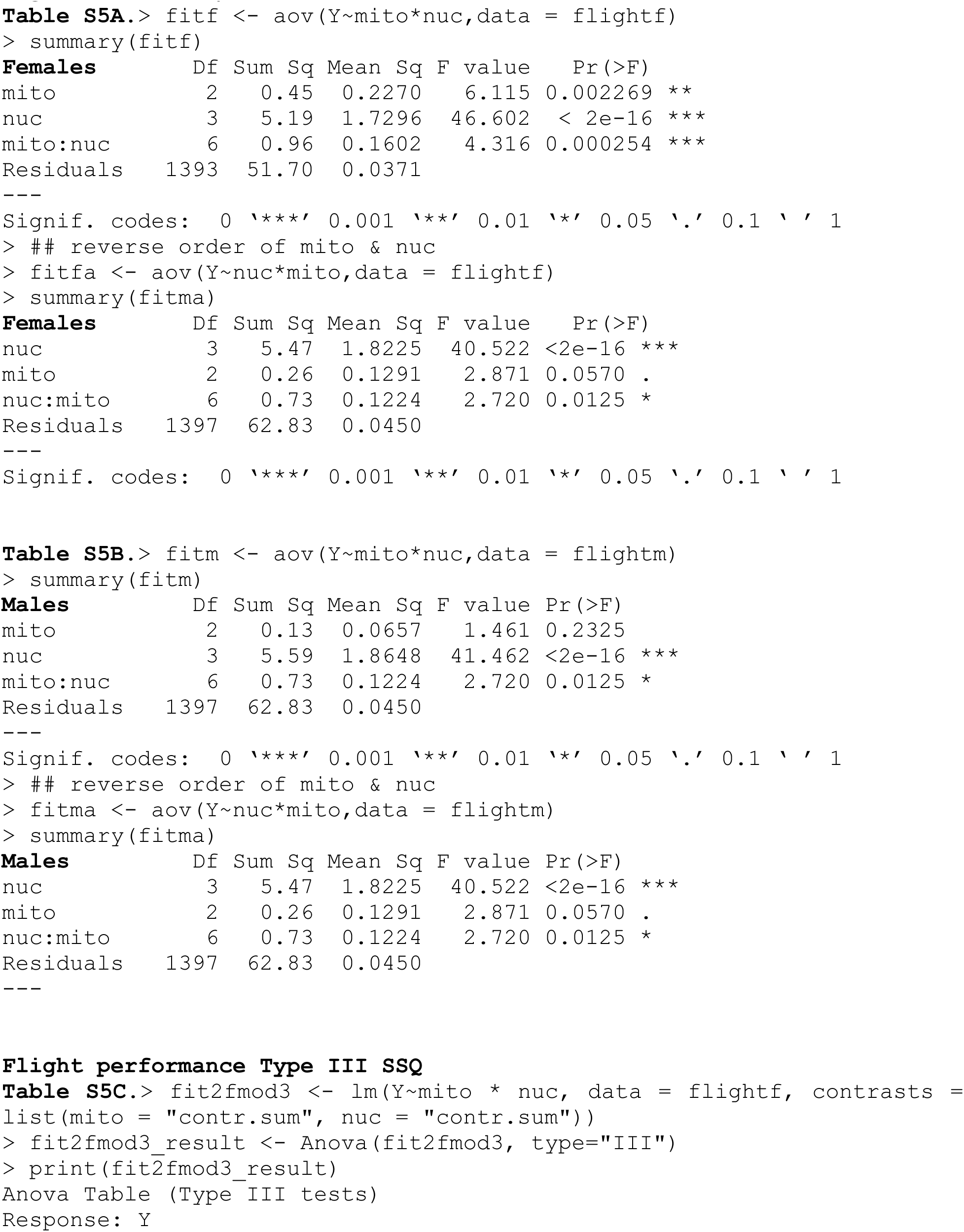
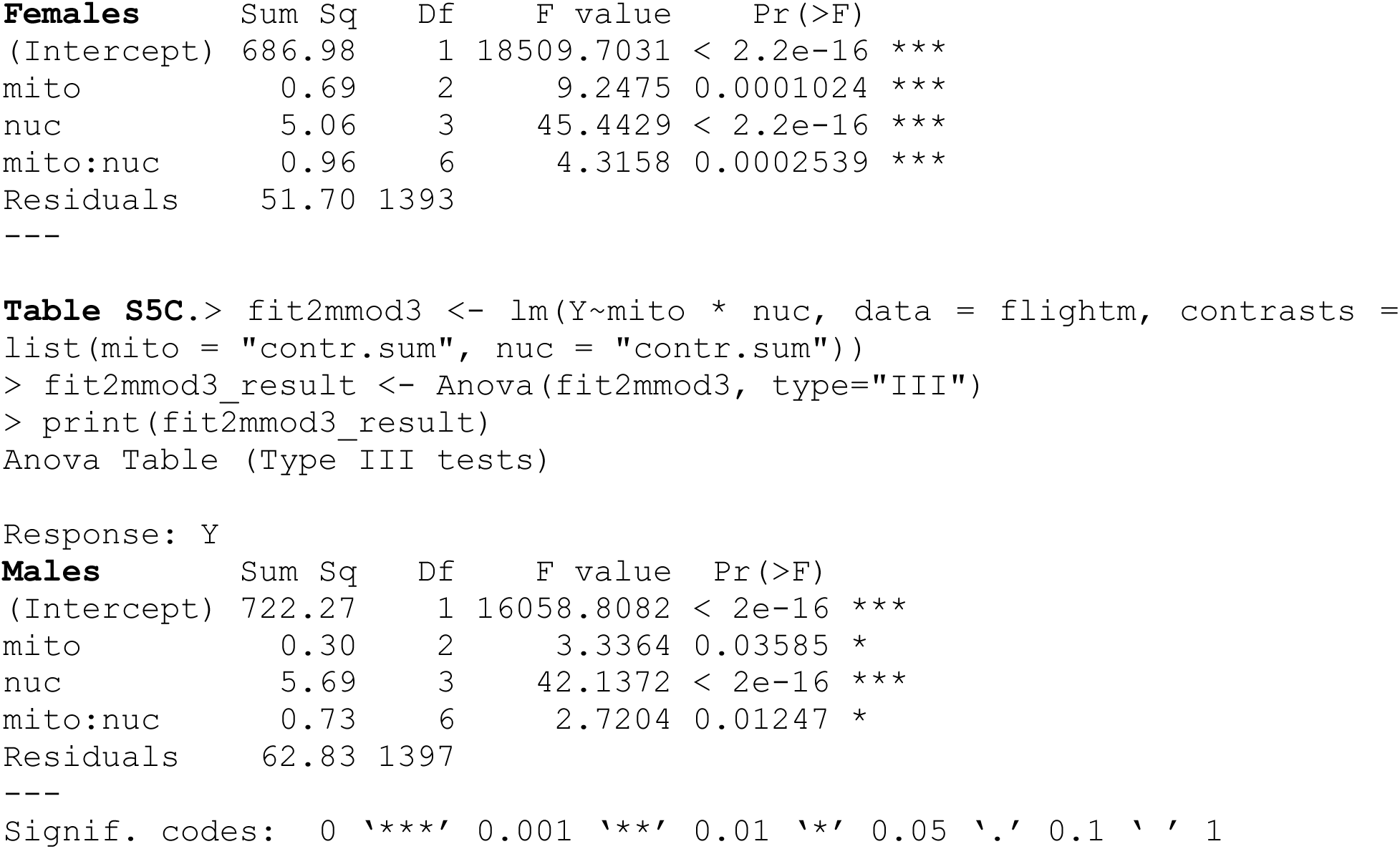
Flight performance (1491, 4959, 7837, *w^1118^*) 3 × 4 data set.

**Table S6.**
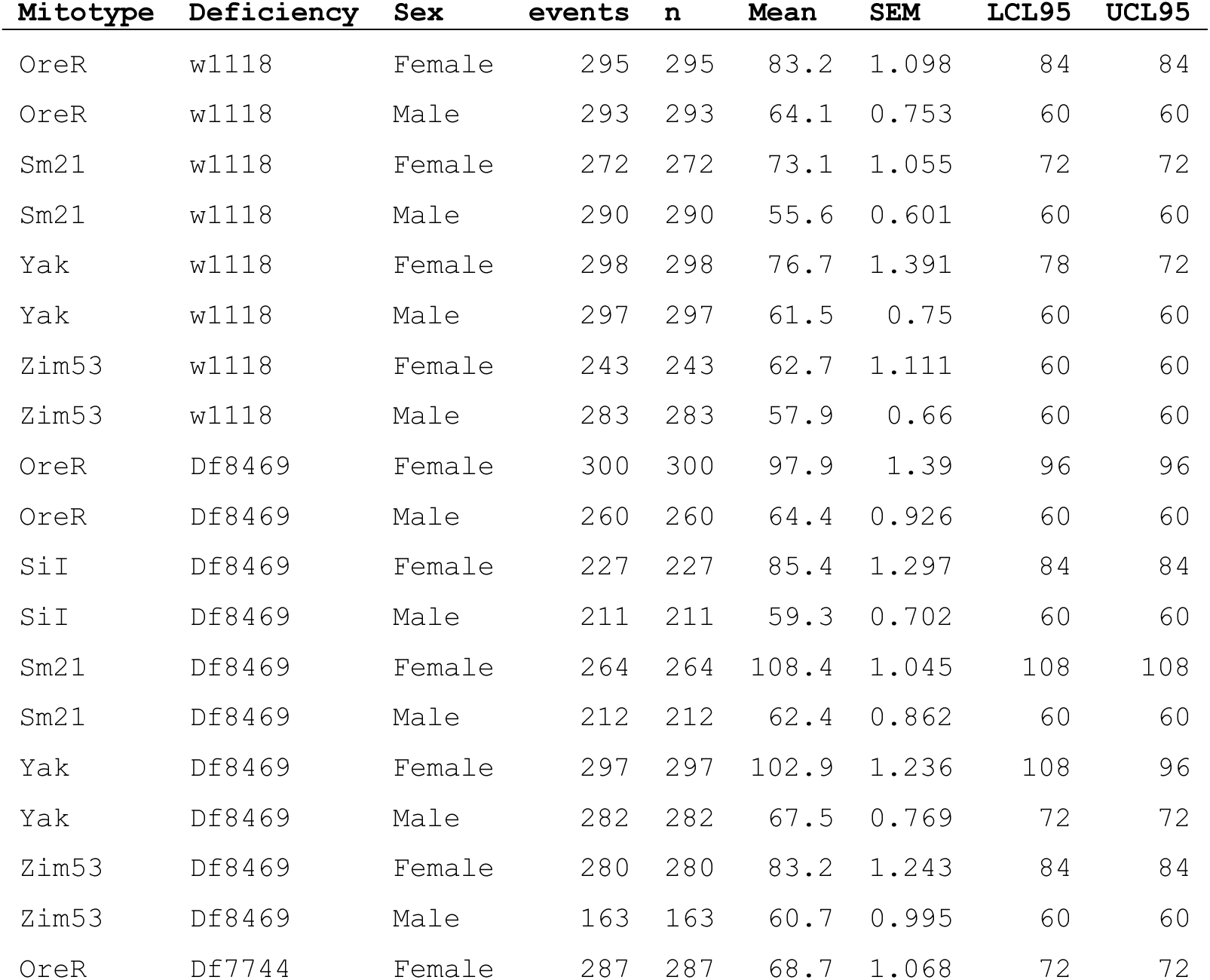
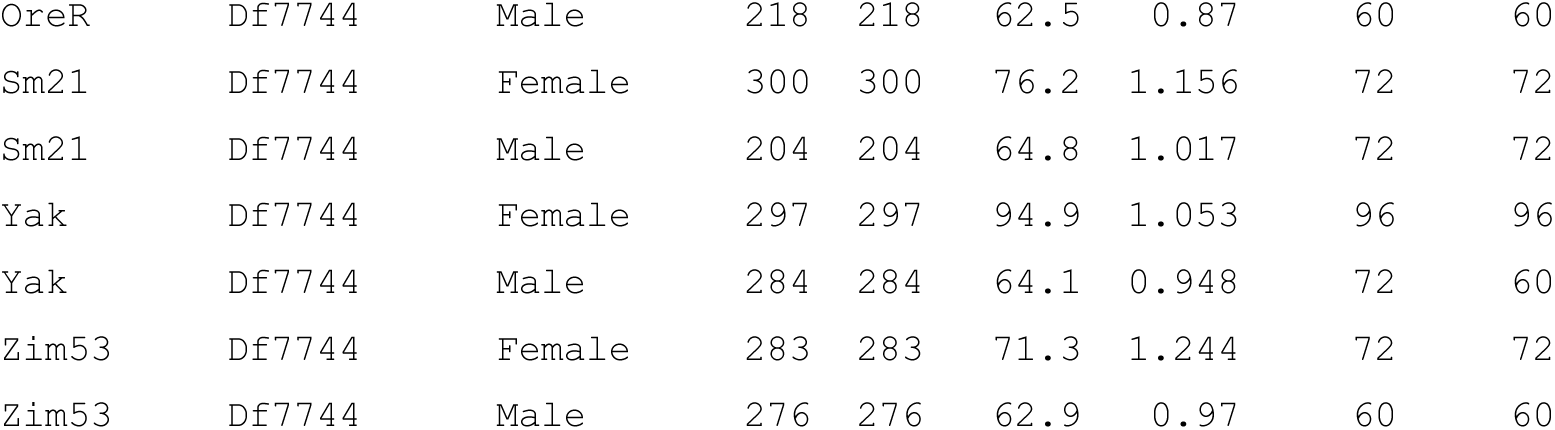
Statistical analyses of survival under starvation.

